# Reversible silencing of endogenous receptors in intact brain tissue using two-photon pharmacology

**DOI:** 10.1101/515288

**Authors:** Silvia Pittolo, Hyojung Lee, Anna Lladó, Sébastien Tosi, Miquel Bosch, Lídia Bardia, Xavier Gómez-Santacana, Amadeu Llebaria, Eduardo Soriano, Julien Colombelli, Kira E. Poskanzer, Gertrudis Perea, Pau Gorostiza

## Abstract

The physiological activity of proteins is often studied with loss-of-function genetic approaches, but the corresponding phenotypes develop slowly and can be confounding. Photopharmacology allows direct, fast and reversible control of endogenous protein activity, with spatiotemporal resolution set by the illumination method. Here, we combine a photoswitchable allosteric modulator (alloswitch) and two-photon excitation (2PE) using pulsed near-infrared lasers to reversibly silence mGlu_5_ receptor activity in intact brain tissue. Endogenous receptors can be photoactivated in neurons and astrocytes with pharmacological selectivity and with an axial resolution between 5 and 10 μm. Thus, two-photon pharmacology (2PP) using alloswitch allows investigating mGlu_5_-dependent processes in wild type animals, including synaptic formation and plasticity, and signaling pathways from intracellular organelles.

## Introduction

Understanding the physiological function of a gene requires pinpointing the exact time course, spatial distribution, and functioning of the encoded protein, from cells to living organisms. This challenge is tackled with techniques based largely on gene targeting and pharmacology. Genetic methods transiently or permanently remove the target protein (knock-out,^1^; knock-down,^2^), thus allowing to spot the effects of protein loss in a genetically-modified organism, but the actual control over the target is intrinsically limited to a timescale of hours to days *(i.e.* the time course of protein expression), and to the cellular specificity of the available molecular markers. Pharmacological blocking of receptor protein function does not reproduce accurately the complex physiology of receptors because spatially-restricted or dynamic control over their targets cannot be exerted. However, recently reported light-regulated drugs (photopharmacology or optopharmacology) can mimic complex receptor kinetics^4^ and dynamics of cell-to-cell communication in intact brain tissue using patterns of illumination^5^. Light drives reversible photoisomerization of such drugs from their active to inactive form and back, and allows turning protein function on and off at illuminated locations and timescales that match the kinetics of the receptor, such that the spatiotemporal resolution for protein switching is only limited by the illumination method.

Photopharmacology can be used in wild type organisms to acutely block the target protein widely, and then “restore” it at any combination of locations and times defined by the illumination pattern. Thus, it allows scrutinizing the role of a protein from the consequences of its presence, instead of its absence, making this technique emerge as an alternative to classical pharmacology (which is limited by the timing required for drug application, diffusion and removal^7^) as well as to genetic targeting (by avoiding physiological compensations and off-target effects often observed with these approaches^6^). The reported spatiotemporal precision of photopharmacology in manipulating neural circuits is comparable to that of optogenetics^8,9^, without the possible drawbacks of distorting cell physiology by overexpression and immunogenicity of microbial proteins^10,11^.

One major limitation of all current photopharmacological approaches is their application in three-dimensional tissues. Our and other groups have developed over the past years a set of freely-diffusible photopharmacological compounds that orthosterically or allosterically target different neurotransmitter receptors (ionotropic [^12–15^] and metabotropic glutamate receptors[^16,17^], μ-opioid receptors[^18,19^], GABA_A_ receptors [^20^], nicotinic ACh receptors [^21^], adenosine receptors [^22^]) and could be used to tackle important research questions in the field, but in practice none of the related published works clearly demonstrates the potential of this technique for activating neurotransmitter receptors with axial specificity.

The majority of these studies show that these molecules can be selectively turned on and off using illumination by single-photon excitation (1 PE), which causes photon absorption throughout the optical axis, and thus photopharmacological drugs are photoisomerized in a large number of neurons at a time. The photostimulated volume can be reduced using pulsed illumination with femtosecond near-infrared (NIR) lasers^23^, which enable two-photon excitation (2PE) of photosensitive molecules to be constrained to physiologically relevant dimensions, such as dendritic spines or individual cells in a circuit^24^. Despite advances in 2PE of opsins^25^, the axial selectivity that can be achieved using photopharmacology and 2PE is still unknown^15,26,27^.

In order to photocontrol the activity of a specific type of receptor with three-dimensional resolution in a physiological context (e.g. in the brain), it is necessary to take a photoswitchable ligand having the highest pharmacological selectivity. Currently, the best available option is alloswitch and its analog compounds, which are freely-diffusible allosteric modulators of metabotropic glutamate 5 (mGlu_5_) receptors^16,28^. These receptors are important modulators of neurotransmission and synaptic plasticity^29^, and are involved in several neurological conditions^30,31,32,33,34,35^. Alloswitch and its analogs are subtype-selective, allosteric ligands with nanomolar activity that can turn on and off endogenous mGlu_5_ receptors *in vivo.* They exist in a thermodynamically-stable, inhibitory *trans* isomer that prevents receptor activation, and contain a light-sensitive moiety based on azobenzene (phenylazopyridine) that can be photoisomerized to its non-inhibitory *cis* configuration using 1PE at visible wavelengths.

Here, we demonstrate that alloswitch can be efficiently photoisomerized using 2PE, we determine the axial resolution of this method, and we establish its feasibility in cultured cells and acute brain slices. This way, the pharmacological blockade of mGlu_5_ is released exclusively using light, which offers an opportunity to silence mGlu_5_ receptors widely, and then photoactivate them at selected locations and times. The rapid and reversible silencing of neurotransmitter receptors in acute brain slices by 2PE of an allosteric photoswitch offers new opportunities to study neuromodulation in intact neuronal circuits and 3D tissues with unprecedented pharmacological selectivity, tissue depth and spatial resolution, and will be invaluable to understand how these receptors function in the unmodified brain.

## Results

### mGlu_5_ receptors are photocontrolled by pulsed light in “all-2PE” experiments

To monitor and manipulate the activity of mGlu_5_ receptors with 2PE of the freely-diffusible drug alloswitch (**Figure 1A**), we used an inverted laser-scanning microscope equipped with a pulsed NIR laser (see **Methods 2** for details). For the initial characterization, we overexpressed the mGlu_5_ receptor fused to the fluorescent reporter eYFP in HEK cells (**Figure 1B**). The activity of mGlu_5_ can be monitored with fluorescence imaging of intracellular Ca^2+^ using chemical dyes and genetically encoded sensors, both at 1PE and 2PE. The expected wavelength for alloswitch isomerization at 2PE is around 780 nm (twice the optimal wavelength for 1PE, 390 nm;^16^. Since switching the power of our NIR laser was faster than switching its wavelength during 2PE imaging and activation, we decided to monitor Ca^2+^ at 780 nm and low excitation intensity using the fluorescent Ca^2+^ indicator Fura-2, which has a good absorption under pulsed NIR light^36^ and no spectral overlap with the eYFP reporter (see **Figure S1 and Methods 3.3.3** regarding single-wavelength imaging with Fura-2). To photocontrol mGlu_5_ activity, we bath-applied alloswitch along with an mGlu_5_ agonist (quisqualate) to generate a ‘ready-steady-go’ state in the cells (**Figure 1A**), whereby, in the absence of illumination, the allosteric antagonist *trans*-alloswitch would put a brake to the intracellular Ca^2+^ events initiated by the mGlu_5_ agonist, whereas its photoisomerization to the *cis* (non-inhibiting) isomer would prompt the onset of Ca^2+^ activity. After mGlu_5_ inhibition was confirmed by silencing of responses to agonist (**Figure S1B**), we used a minimal laser power (3 mW, 780 nm) to visualize Ca^2+^ by 2PE-imaging of Fura-2, and an increased power (12 mW) to produce 2PE-isomerization of alloswitch to the non-inhibiting *cis*-isomer, leading to cell activation (see strategy for achieving interleaved 2PE-imaging and 2PE-stimulation, namely “all-2P experiments”, in **Figure S1C** and **Methods 3.3**; see **Figure S1D–E** about increased fluorescence of sample during photostimulation). Laser power was kept to minimal values for 2PE-imaging to minimize unwanted 2PE of alloswitch molecules due to imaging of Fura-2.

**Figure 1:**
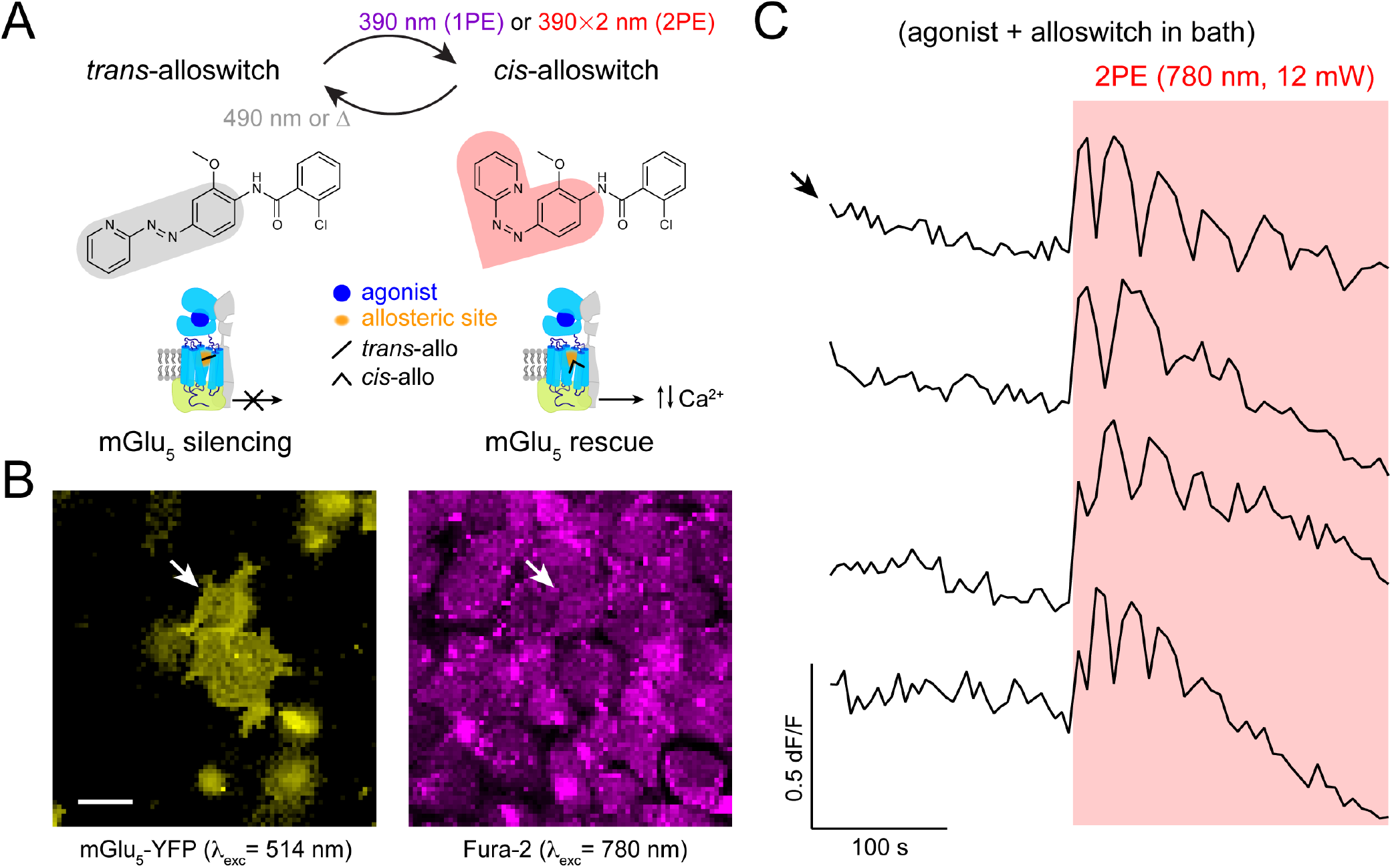
Two-photon excitation of alloswitch at NIR wavelengths in cultured cells. **A Top**: *Trans*→*cis* photoisomerization of alloswitch occurs by single-photon excitation (1PE) under violet light (390 nm). Using pulsed-lasers for two-photon excitation (2PE), the expected wavelength for photoswitching is double that required for 1 PE. Back-isomerization occurs by thermal relaxation (Δ; half-life of *cis* isomer, t½ = 116 s;^16^ or under 490 nm light. The photosensitive core of alloswitch – azobenzene – is highlighted in its *trans* (grey shade) and *cis* configurations (red shade). Bottom: *Trans*-alloswitch silences active, agonist-bound mGlu_5_ receptors by binding at the allosteric pocket, whereas *trans*→*cis* photoisomerization of alloswitch rescues the silenced, agonist-bound receptors at the site of illumination by releasing intracellular agonist-induced Ca^2+^ oscillations. **B** Cultured HEK cells expressing mGlu_5_-YFP (yellow, left) and loaded with Fura-2 (magenta, right) for two-photon imaging of Ca^2+^ (760–820 nm, shown here is 780 nm). Arrows indicate the cell whose Ca^2+^ photoresponse is shown in the top trace of panel **C**. Scale bar is 20 μm. **C**. Representative traces of Ca^2+^ oscillations induced by 2PE of alloswitch (1 μM) with quisqualate (mGlu_5_ orthosteric agonist, 3 μM) in bath (arrow indicates trace of cell shown in **B**). Ca^2+^ oscillations were elicited by 2PE of alloswitch at 780 nm (red box; 12 mW scans, see ***Methods section*** and **Figure S1C** for details on illumination patterns) and are depicted as fluorescence changes of Fura-2 relative to baseline (dF/F; excited at 780 nm, 3 mW scan; see **Figure S1A** for direction of fluorescence changes of Fura-2 excited at 780 nm). See similar experiments for other alloswitch analogs in **Figure S3**.

We observed that 2PE raster scans at 780 nm, 12 mW and 10 μs dwell time (see **Methods 2 and 3.3**) were sufficient to elicit mGlu_5_-dependent intracellular Ca^2+^ oscillations, indicating efficient alloswitch photoisomerization and release of the ‘brake’ exerted by alloswitch on the receptors (**Figure 1C**). Note that all cells displaying Ca^2+^ oscillations under 2PE were eYFP^+^, which indicates that responses were mediated by mGlu_5_ and rules out artifacts due to photostimulation alone.

To determine the optimal wavelength for 2PE of alloswitch, we repeated “all-2PE” experiments at different spectral positions in the NIR range between 760 and 820 nm. Fura-2 allows 2PE fluorescence imaging of mGlu_5_-dependent Ca^2+^ oscillations at all wavelengths in that range (**Figure S1A**), and can be combined with 2PE-isomerization of alloswitch at the same wavelengths (**Figures 2, S1C** and **S2**). We found that 2PE at 780 nm produced the optimal photoisomerization of alloswitch. Although the maximum oscillatory frequency was lower at 780 nm than at 760 nm (**Figure 2A**), oscillations at 780 nm lasted longer (**Figure 2B**), were induced faster (**Figure 2C**) and achieved more oscillatory events (**Figure 2D**), despite a slightly lower laser power than that measured at 800 nm (**Figure S2E**).

**Figure 2:**
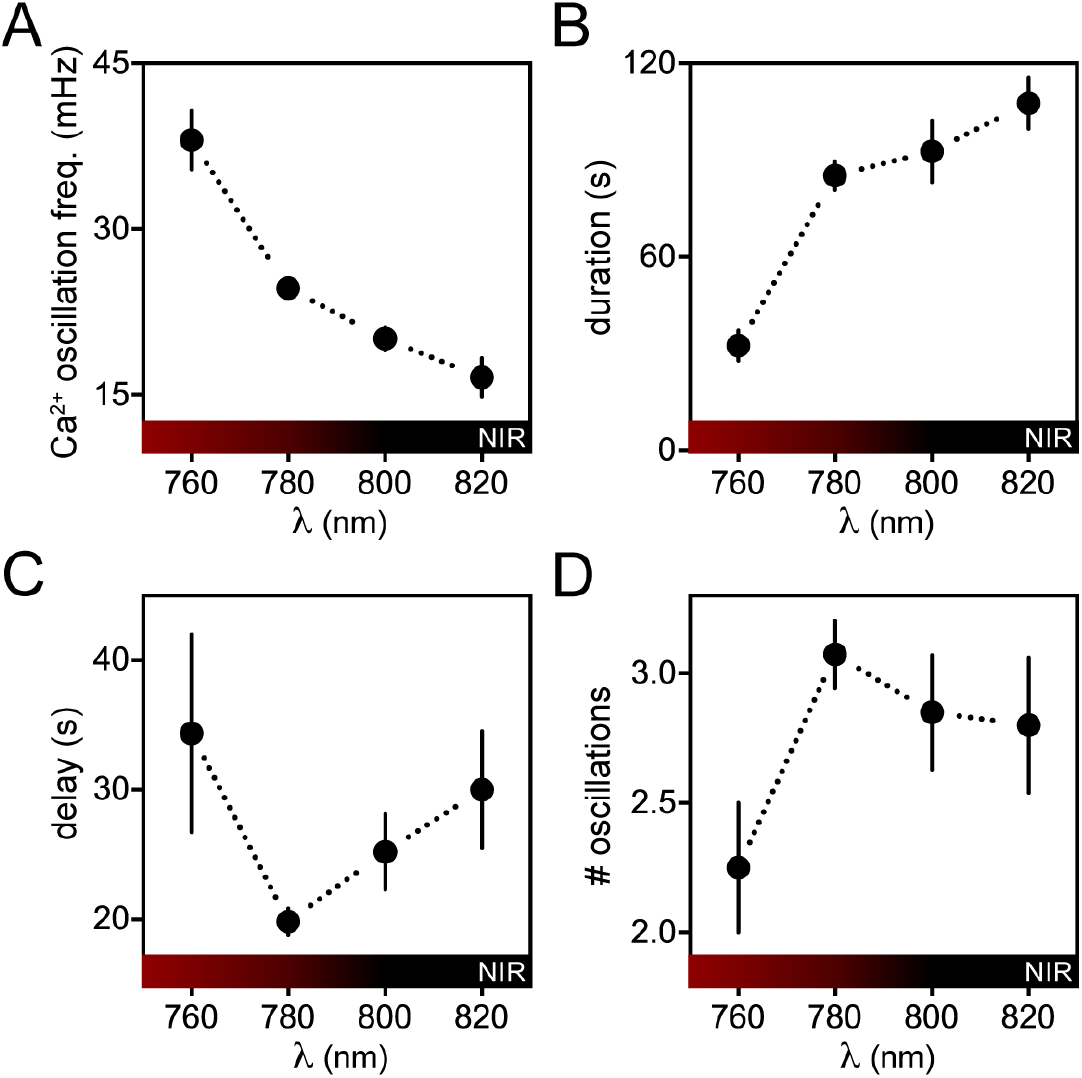
Characterization of the efficacy of responses to 2PE of alloswitch as a function of the pulsed laser wavelength. Experiments were conducted as in **Figure 1C**, in cultured cells expressing mGlu_5_-eYFP and loaded with Fura-2 for 2P-Ca^2+^ imaging; cells were supplemented with agonist (quisqualate, 3 μM) and alloswitch (1 μM) prior to experiments. Wavelength-dependence of frequency of Ca^2+^ oscillations (**A**), duration of the oscillatory behavior (**B**), time for onset of first peak after 2PE initiation (delay, **C**), and average number of oscillations observed during 2PE (# oscillations, **D**). Frequency of calcium oscillations was higher after 2PE at 760 nm compared to other wavelengths (38.1 ± 2.6 mHz, *p* < 0. 05) and at 780 nm compared to 820 nm (24.7 ± 0.7 Hz, *p* < 0.001). Duration of oscillatory response was lower for 760 nm compared to other wavelengths (32.4 ± 4.7 s, *p* < 0.05). Data represent the mean ± s.e.m., n = 4-95 cells; Dunn’s multiple comparison test after the Kruskal-Wallis test, see **Figure S2** for full statistic comparisons.

Several alloswitch analogs have been recently reported that display a variety of pharmacological and photophysical properties^28^, and we selected a few in order to test their ability to photocontrol mGlu_5_ receptors at 2PE. We repeated the previous experiments using analogs bearing substitutions at the distal phenyl ring, and found that all of them were responsive at similar wavelengths (**Figure S3**).

### Efficacy of 2PE and 1PE to rescue silenced mGlu_5_ receptors using alloswitch in HEK cells

2PE of molecules using pulsed lasers requires the coincident absorption of two photons and therefore occurs with lower probability than 1PE and mainly at the focal volume^23^. In the case of photoswitches, this translates into generally lower isomerization efficacy at 2PE than at 1PE even if photoisomerization quantum yields are unchanged^27^. We set out to compare the efficacy of 2PE versus 1PE of alloswitch (**Figure 3A**) by measuring responses repetitively and reproducibly in the same cell to avoid variability due to heterogeneity of receptor density and localization. For this purpose, we used the genetically encoded Ca^2+^ sensor GCaMP6s^37^, whose excitation wavelength does not cross-excite alloswitch and allows longer experiments than Fura-2 without risk of photostimulation due to imaging. GCaMP6s was co-expressed in HEK cells with mGlu_5_ receptors lacking the eYFP tag, to avoid fluorescence crosstalk (**Figure 3B** and corresponding **Movie S1**).

**Figure 3:**
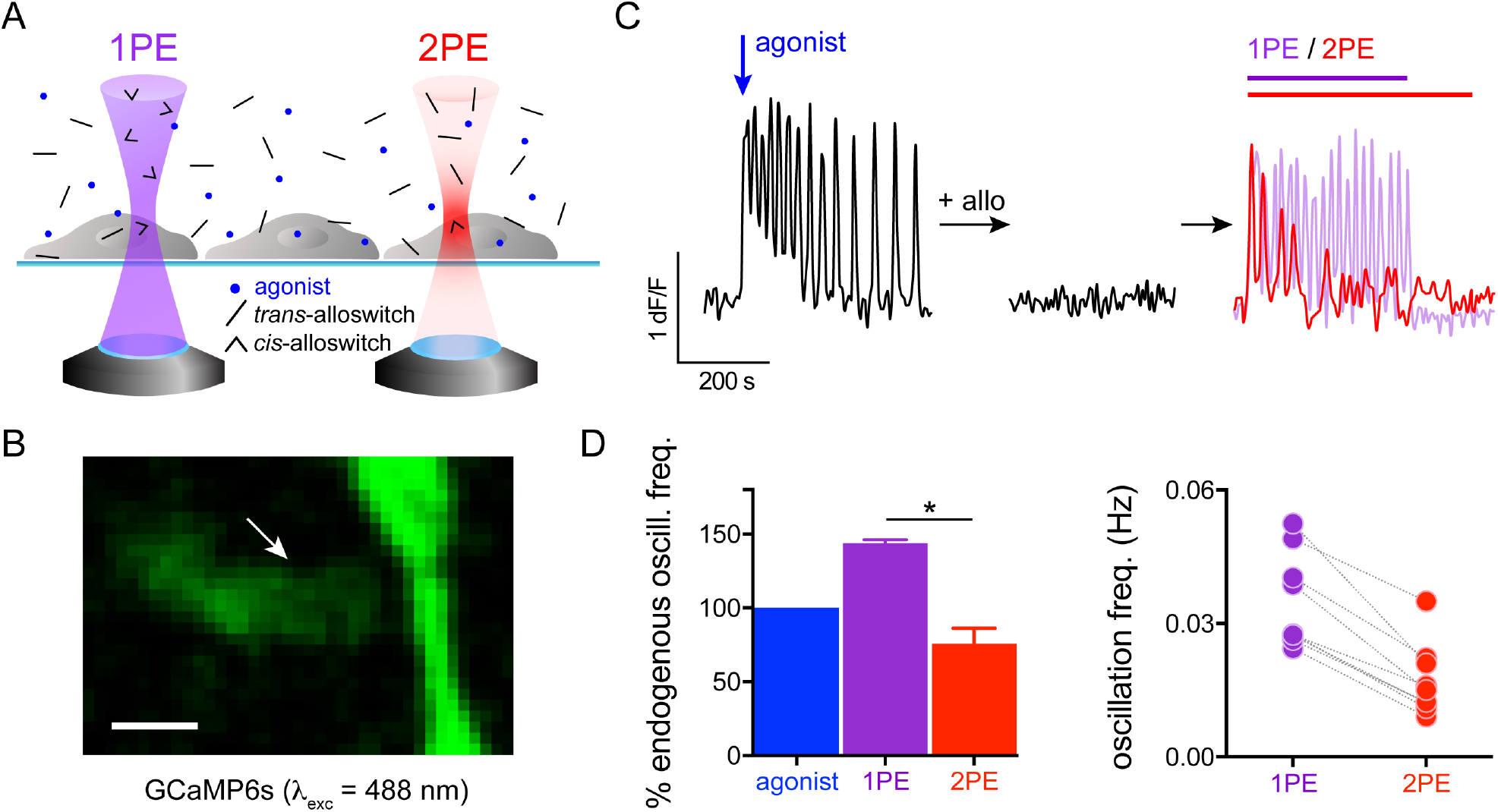
Efficacy of mGlu_5_ rescue by 1PE or 2PE of alloswitch in cultured cells. **A** Principle of alloswitch photoactivation *(trans:* black sticks; *cis*: arrowheads) using continuous-wave (CWL) or pulsed lasers (PL). Light-cones represent probability of efficient excitation with one (1 PE, 8violet) or two coincident photons (2PE, red). Cells expressing mGlu_5_ are cultured on a glass coverslip and supplemented with agonist (blue dots) and alloswitch, and light from the objective of an inverted confocal microscope is focused at the cell plane. Light from a CWL excites molecules in solution at 1PE along the entire light cone (note *cis*-alloswitch all along the light cone), whereas light from a PL excites molecules (2PE) within its point spread function (PSF) volume (for most 2PE systems, 1 μm^3^; for representation, 2PE spot is exaggerated compared to cell size). **B** The genetically-encoded Ca^2+^ sensor GCaMP6s (fluorescence excited at 488 nm) was co-expressed with mGlu_5_ in HEK cells. Arrow indicates cell in **C** (see also **Movie S1**). Scale bar is 10 μm. **C** Representative Ca^2+^ trace (expressed as changes in GCaMP6s fluorescence, dF/F) from the cell indicated by arrow in **B** in different conditions. **Left**, response of the untreated cell elicited by addition of an mGlu_5_ agonist (quisqualate, 3 μM, blue arrow); right, the same cell after treatment with alloswitch (1 μM) and photoactivated at single-photon (1PE, violet line and trace) or two-photon (2PE, red line and trace). See **Methods 3.4** for GCaMP6s imaging conditions and **Methods 5.2** for alloswitch photostimulation using 1PE and 2PE. **D** Quantification of oscillatory Ca^2+^ events from experiments as in **C. Left**: Percentage of Ca^2+^ oscillations frequency elicited by receptor activation with 1PE (violet bar, 144±2%) or 2PE (red bar, 76±11%) of alloswitch, relative to the oscillatory frequency generated by agonist alone acting on the endogenous mGlu_5_ receptors. Data are mean ± s.e.m., n = 4 cells. * *p* < 0.05, Dunn’s multiple comparison test after the Friedman test for matched data. **Right**: 2PE of alloswitch generated on average 48 ± 4% of the frequency of Ca^2+^ oscillations induced by 1PE in the same cell. Data are mean ± s.e.m., n = 9 cells from two independent experiments. Paired t-test, *p* < 0.0001.

We activated mGlu_5_ receptors by adding to the bath the agonist quisqualate (3 μM) and recorded the time course of Ca^2+^ responses (**Figure 3C left**). In cells displaying oscillatory behavior, the frequency of Ca^2+^ oscillation is proportional to the number of available mGlu_5_ receptors^38^. We then silenced mGlu_5_ responses by addition of 1 μM alloswitch in the bath (**Figure 3C middle**), and sequentially photostimulated cells in the field of view at 1PE and 2PE (**Figure 3C right**). We compared the efficacy of alloswitch to photoinduce mGlu_5_-dependent Ca^2+^ oscillations at 1PE or 2PE relative to the endogenous activity of the receptor (expressed as oscillatory frequency of cells in response to agonist alone, **Figure 3D left**), and we found that 2PE produced oscillatory response frequencies that were 76 ± 11 % of those induced by the agonist, and 53% of those induced by 1PE. Note that the oscillatory activity in response to 1PE of alloswitch was higher than in response to the applied agonist alone, as previously reported^16^. Individual cells displaying oscillatory responses to both 1PE and 2PE allowed us to compare the fraction of mGlu_5_ receptors photoactivated by 1PE and 2PE, being the number of expressed receptors and signal transduction machinery unchanged for a given cell within the time course of our experiments. We found that the frequency of oscillations with 2PE of alloswitch was 48 ± 4% of that obtained with 1PE in the same cell (**Figure 3D right**), which suggests that about half the receptors are recruited in the small 2PE volume compared to 1PE of the entire cell.

### Silenced receptors are rescued by 2PE of alloswitch with axial-plane selectivity in a cell monolayer

An advantage of 2PE over 1PE is that the former reduces out-of-focus excitation in the axial direction by orders of magnitude, restricting to micrometric volumes the excitation of molecules^23^. This is well established in the case of fluorescent molecules, but it remains unexplored for photoswitchable compounds^15,26,27^. Here, we asked whether alloswitch could be photoisomerized to its non-antagonizing *cis* isomer with axial precision, despite diffusion of the inhibiting *trans* isomer from outside the 2PE volume. To answer this question, we co-expressed mGlu_5_ and GCaMP6s in HEK cells and repeated 1PE and 2PE at various axial distances above the cell focal plane (**Figure 4A-B** and corresponding Movie S2). Cell responses to the receptor agonist were recorded, then alloswitch was applied to the bath to inhibit the induced Ca^2+^ oscillations (**Figure 4C left**). 1PE of alloswitch photoactivated cells regardless of the axial position of photostimulation, with the maximal distance tested 30 μm above cells (**Figure 4C**).

**Figure 4:**
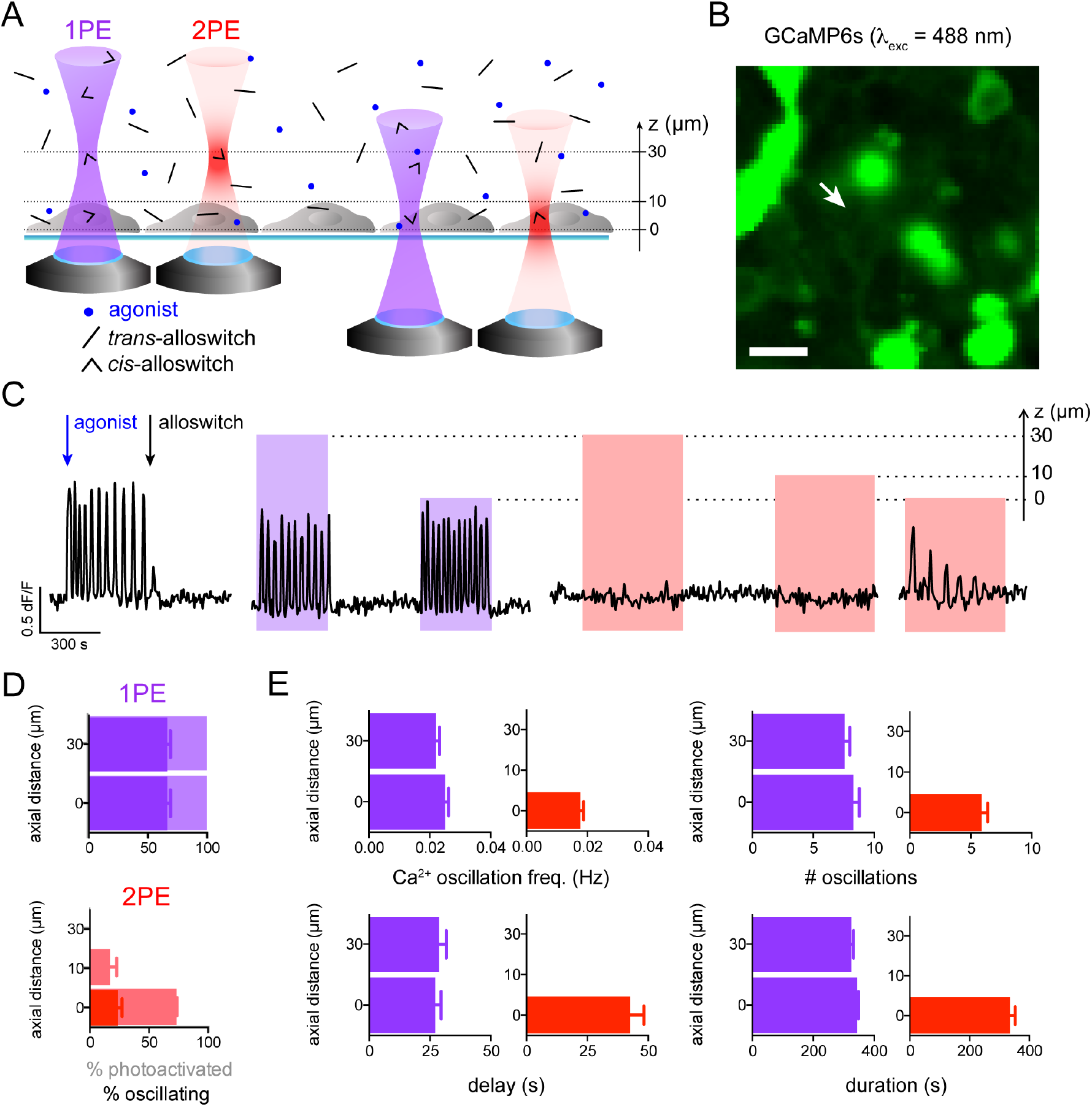
2PE of alloswitch in free-diffusion medium enables axially-resolved photoactivation of mGlu_5_. **A** Experimental setup. Cells expressing mGlu_5_ were seeded on a glass coverslip, and light from the objective of an inverted microscope was focused at different axial distances (z = 30, 10 or 0 μm) upwards from the cell focal plane. 1PE of alloswitch occurs at any axial distance, whereas 2PE excites alloswitch in the proximity of the sample only when focused at the cell plane (z = 0). **B** Fluorescence image of HEK cells co-expressing mGlu_5_ and GCaMP6s for Ca^2+^ imaging. Arrow indicates cell recorded in **C**. Scale bar is 20 μm. **C** Control of Ca^2+^ responses in the axial direction using 2PE of bath-applied alloswitch in a representative cell (indicated by arrow in **B**). Photoactivation of cell responses occurs regardless of the focal plane using 1PE of alloswitch, and is focal-plane (z = 0) selective with 2PE. Ca^2+^ oscillations were elicited by addition of an mGlu_5_ agonist (quisqualate, 3 μM, blue arrow) and blocked by bath-application of alloswitch (1 μm, black arrow). Light for alloswitch excitation is focused at different axial distances above the Ca^2+^-imaging plane, as indicated by the right axis (z, μm). See **Methods 3.4** and **5.2** for imaging and stimulation conditions. **D** Comparison of the cell responses using 1PE (top) or 2PE (bottom) at different axial distances above cells, measured as % of photoactive cells (positive Ca^2+^ changes; faint colors) and % of cells showing oscillatory behavior (solid colors). All cells responded to 1PE regardless of the focal plane, and 66 ± 3% of them displayed an oscillatory behavior. With 2PE at the focal plane (0 μm) photoactive cells were 73.0 ± 0.3%, and oscillating cells were 24 ± 4% (36% of the cells found to oscillate at 1PE). When 2PE was applied at 10 μm above the focal plane, 17 ± 6% cells were photoactive. 2PE induced no oscillations when performed at axial distances of 10 or 30 μm. Percentages were calculated relative to the total number of cells found to respond to 1PE at the focal plane (0 μm). n = 67 cells from 2 independent experiments. See full dataset in **Figure S4E.** **E** The oscillatory behavior of cells photoactivated at 1PE or 2PE and different axial distances was quantified by measuring the frequency and total number of calcium oscillations, time at which the first oscillation occurred after begin of illumination (delay), and overall duration of the oscillatory behavior (duration). The frequency of Ca^2+^-oscillations when photoactivating at 2PE at the focal plane was lower than at 1PE (0.018 ± 0.001 Hz and 0.025 ± 0.001 Hz respectively, *p* < 0.01). The maximum number of oscillations achieved at 2PE at the focal plane was also lower than at 1PE (8.3 ± 0.5, *p*<0.05). The delay for response onset with photostimulation at the focal plane was longer in trend for 2PE than 1PE (42 ± 6 and 27 ± 3 s, p=0.25). The duration of oscillatory behavior was unaffected by light stimulation. Cell responses did not change for 1PE at either axial distance. Data are mean ± s.e.m; statistic test used was the Kruskal-Wallis test with Dunn’s correction. 1PE, n = 45 from 2 independent transfections; 2PE, n = 27 from 3 independent experiments. See full dataset and statistics in **Figure S4F**.

Of the total number of photoresponsive cells (displaying changes in intracellular Ca^2+^, see **Methods** for classification details and **Figure S4C-D** for example traces; faint colors in **Figure 4D top**), 66 ± 3 % displayed oscillatory responses at both 0 and 30 μm (dark bars in **Figure 4D top**; see also **Figure S4E**). About 75% of cells that responded to 1PE were also photoactivated by 2PE at the cell plane (0 μm), and 24 ± 4% displayed oscillatory responses (**Figure 4D bottom**; see **Figure S4C–D** for example traces). In contrast, photoisomerizing alloswitch with 2PE only 10 μm above the cell plane produced responses in only 17 ± 6% of cells, and none when the photostimulation plane was 30 μm above the cell focal plane (**Figure 4D bottom** and **Figure S4E**). We further quantified the frequency, oscillation number, onset time (delay) and duration of the Ca^2+^ responses, and found that 2PE at the cell plane elicited slightly lower frequencies, fewer oscillations, and longer delay than 1PE, whereas response duration was similar for 2PE and 1PE (**Figure 4E**; see statistics in **Figure S4F**).

Together, these results demonstrate that mGlu_5_ photoactivation in a volume restricted by 2PE of alloswitch is possible despite diffusion of inhibiting *trans* isomers into the 2PE volume. The axial resolution in our experimental conditions allows selective activation of mGlu_5_ signaling within 10 μm of the target receptors. The fact that the basal cell surface in contact with the glass coverslip (0 μm) can be photoactivated more efficiently than the apical side exposed to the bath (10 μm) suggests that more receptors are photorescued at the basal side, probably because its flat shape matches better the 2PE plane than the curved contour of the apical side.

### Alloswitch can be used for functional silencing and light-induced rescue of mGlu_5_ receptors in acute rodent brain slices

To test whether alloswitch can photocontrol endogenous mGlu_5_ receptors expressed by both neurons and astrocytes in intact neural tissue, we set up photoswitching experiments in acute rodent brain slices while monitoring activity in both cell types using the Ca^2+^-sensitive dye OGB-1 (**Figure 5A-B**). We used rat pups (postnatal days 6-15) due to the high expression of mGlu_5_ receptors in neonatal (**Figure S6A**) compared to adults brain tissue^39^, and we targeted hippocampal CA1 region because of the documented mGlu_5_ receptor expression in this brain area^40^. We used a glass micropipette placed on top of the field of view for local delivery of the mGlu_1_/mGlu_5_ agonist DHPG by pressure-ejection (**Figure 5B**). The circulating bath contained LY367385 to antagonize mGlu_1_ responses, and tetrodotoxin (TTX) to prevent propagation of responses to neighboring neurons. Photostimulation of alloswitch and fluorescence imaging of OGB-1 were performed using the inverted confocal microscope setup described above (see **Figure S5A** for illumination paradigm). Because loading of OGB-1 is not cell-type selective, recorded Ca^2+^ traces represent responses from a mixed population of neurons and astrocytes.

**Figure 5:**
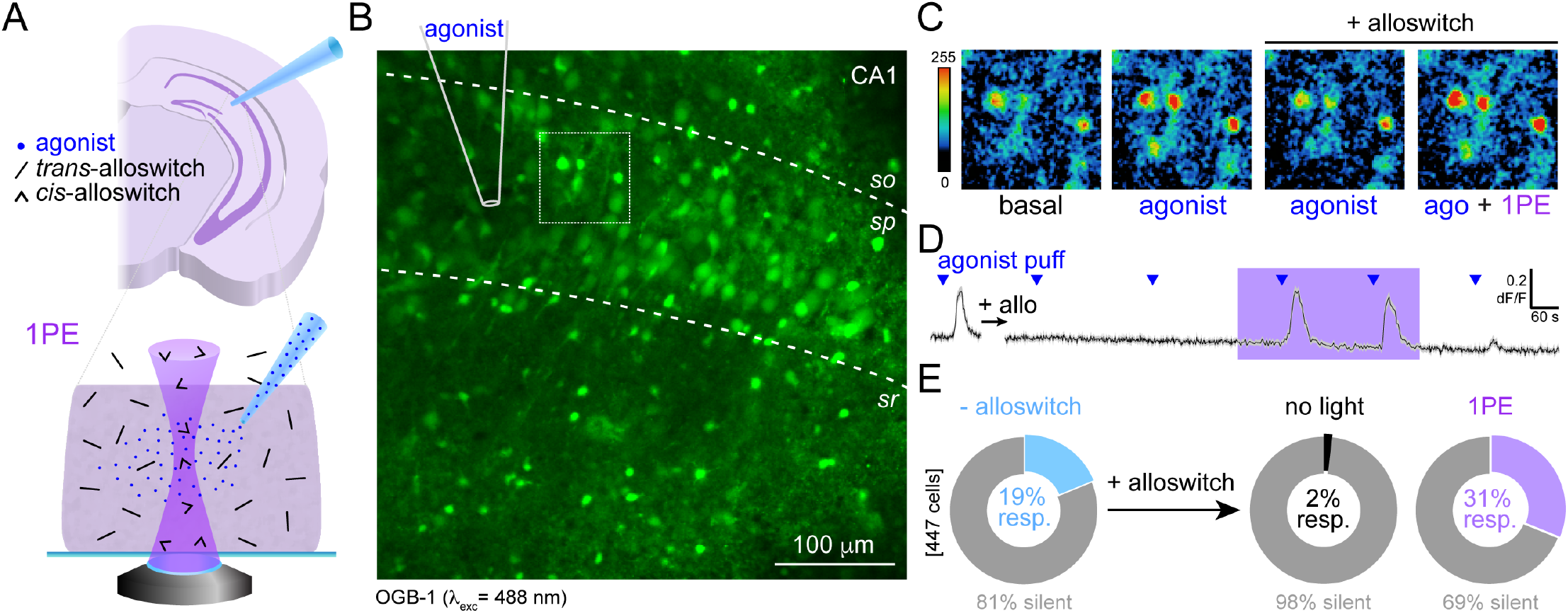
Functional silencing and light-induced rescue of mGlu_5_ receptors in acute rodent brain slices. **A** Experimental design for the activation of mGlu_5_ receptors with light in brain slices (1PE). **Top**: a pipette filled with mGlu_5_ agonist (DHPG, blue dots) is placed on the CA1 region of the caudal hippocampus in a rat brain slice for cell stimulation. **Bottom**: *trans*-alloswitch in bath (which blocks mGlu_5_ responses to agonist) is photoconverted to *cis-* alloswitch (mGlu_5_ block is released) along the light cone of the light source (violet, 1PE). **B** Average fluorescence of OGB-1 loaded on CA1 region of the caudal hippocampus from a recorded slice, indicating strata (dashed lines) and pipette position (solid lines) for local ejection of agonist. Dashed rectangle indicates region magnified in panel C. Scale bar is 50 μm; so, *stratum oriens; sp, stratum pyramidale; sr, stratum radiatum.* Images acquired at 1PE with 488 nm laser line at 2.4 μW power. **C** Magnification of inset in panel **B** in pseudocolors, illustrating the time course of Ca^2+^ responses at time 0 (‘basal’), 40 s after ejection of agonist in absence (‘agonist’) or presence (‘agonist + alloswitch’) of alloswitch in the bath. Response during 1PE of alloswitch is shown 37 s after agonist ejection (‘ago + 1PE + alloswitch’). Scale bar is 10 μm. **D** Average trace of Ca^2+^ responses to agonist ejection at times indicated by blue triangles (1 mM DHPG; 800 hPa, 1 s) acquired before (left) and after (right) addition of alloswitch in bath (‘+ allo’, 10 μM), and during light stimulation in the 1PE regime (violet box, 405 nm laser diode, 58 μW power, 2.5 μs dwell time, and 0.3 Hz illumination rate). Data are mean fluorescence changes of OGB-1 (black) ± s.e.m. (grey) from 39 cells initially responding to agonist. OGB-1 was excited at 488 nm, 2.4 μW, 0.3–1 Hz. See **Figure S5A** for imaging/photostimulation schematics and for control trace without alloswitch. **E** Quantification of the % of cells responding to agonist ejection in experiments as depicted in panel **D**. OGB-1 labeled cells not responding to agonist puffing are shown in grey (silent). Cells responding to agonist before alloswitch treatment were 18.8 ± 0.8% of the total (light blue, ‘-alloswitch’). Adding alloswitch in bath reduced the amount of responding cells to 1.9 ± 1.0% (black, ‘no light’), whereas photoisomerizing alloswitch with 1PE rescued the responses to mGlu_5_-agonist in 31.2 ± 0.2% of the total number of cells (violet, 1 PE, 58–98 μW). The fraction of responding cells is thus significantly reduced by addition of alloswitch (*p*<0.01), and increased by 1PE of alloswitch (*p*<0.05); paired t-test. Percentages are calculated out of OGB-1 labeled cells in a 0.14 mm^2^ area surrounding the pipette. Data are mean ± s.e.m., n = 447 cells from 2 animals.

After establishing Ca^2+^ responses of untreated cells to DHPG ejection (**Figure 5C**, labeled ‘agonist’, compared to OGB-1 fluorescence at rest, ‘basal’), alloswitch was added to the bath (10 μM). The cellular responses mediated by mGlu_5_ were silenced in presence of *trans*-alloswitch (**Figure 5C**, labeled ‘agonist/alloswitch’). Upon raster-scanning the field of view to photoisomerize alloswitch to the *cis*-isomer by 1PE, mGlu_5_ responses to agonist ejection were readily rescued (**Figure 5C**, labeled ‘ago + 1PE/alloswitch’). The time course of OGB-1 fluorescence from an example experiment (**Figure 5D** and Movie S3; see **Figure S5A** for control without alloswitch), and counting of silent and responding cells (**Figure 5E**) show that alloswitch and 1PE enable reversible silencing and light-induced rescue of mGlu_5_ activity in brain slices. DHPG-responding cells drop from 19% to 2% in the presence of 10 μM alloswitch. Interestingly, during 1PE of alloswitch Ca^2+^ responses were recovered in 31 % of the cells in the field of view, a higher fraction (*p*<0.05; paired t-test) than observed prior to alloswitch application (**Figure 5E**).

### Efficacy of 2PE of alloswitch is 30% of 1PE to rescue silenced mGlu_5_ receptors in brain slices

We next aimed to constrain mGlu_5_ activation to the focal plane using 2PE in acute brain slices. We performed Ca^2+^ imaging experiments with LY367385, TTX, and alloswitch in the bath (**Figure 6A**). The same cells were sequentially stimulated with DHPG before, during 2PE and 1PE of alloswitch, and after illumination, as shown in the example traces of **Figure 6B** and Movie S4 (see also **Figure S5** for control experiments without alloswitch). 2PE successfully induced Ca^2+^ responses in 25% of cells in the imaged region compared to 76% of responses with 1PE; in addition, mGlu_5_ events triggered by 2PE displayed 15% lower peak amplitude (**Figure 6C**) and were shorter than 1PE, suggesting a confined photoisomerization of alloswitch and limited number of recruited receptors under 2PE.

**Figure 6:**
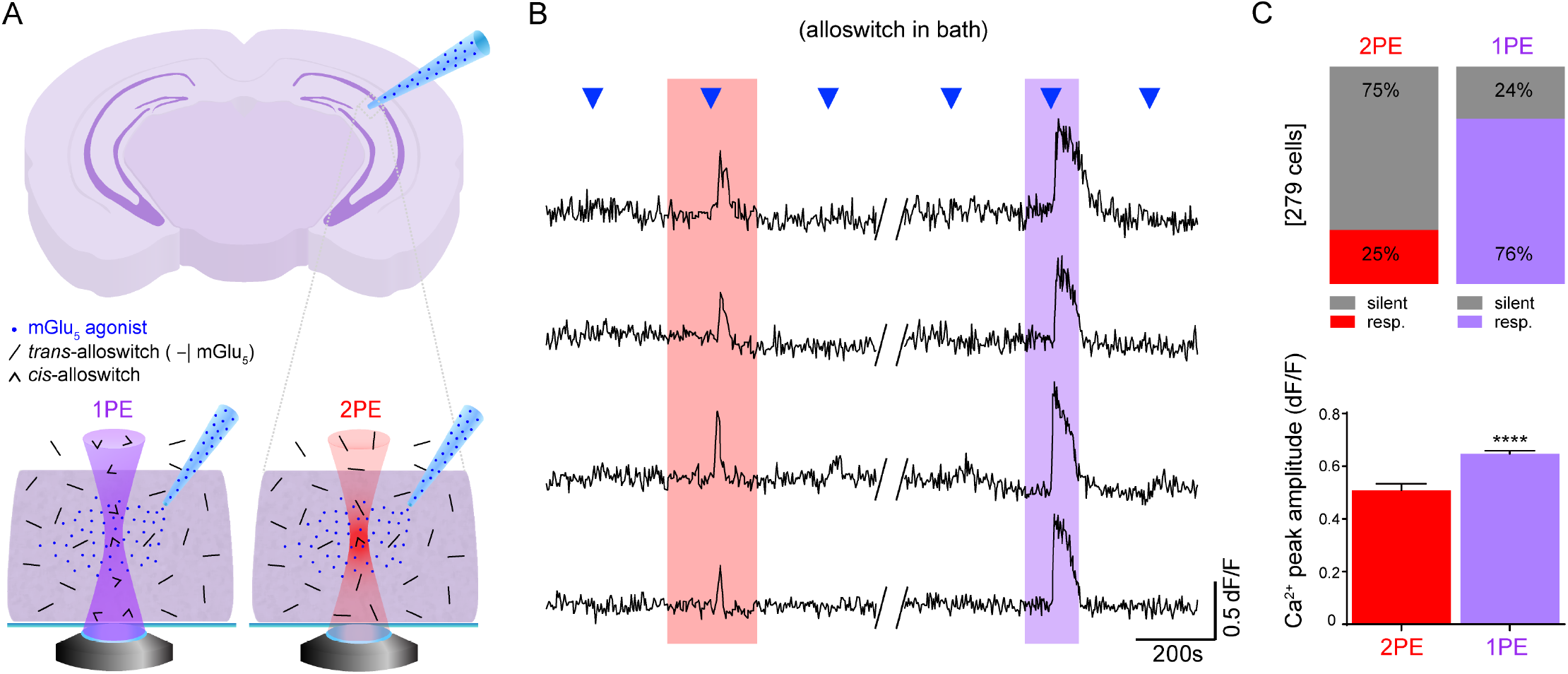
Efficiency of mGlu_5_ activation using 1PE and 2PE of alloswitch in a 3D tissue. **A** Schematics of the experimental design. *Trans*-alloswitch in bath (mGlu_5_-block) is photoconverted to *cis*-alloswitch (mGlu_5_-block is released) along the volume of brain slice illuminated by the light cone of 1PE (violet) or in the focal spot of 2PE (red). Pipette for delivery of mGlu_5_ agonist (DHPG) is placed on top imaged cells in the CA1 region of the hippocampus. **B** Example traces of 4 recorded cells with alloswitch in bath, comparing the Ca^2+^ responses to agonist puffing (blue triangles) before, during, and after 2PE (left, red box) or 1PE (right, violet box) of alloswitch. 1PE and 2PE were achieved with 2 (1PE) or 3 (2PE) consecutive raster scans at a frequency of 0.4 Hz and 0.26 Hz, respectively, with pixel dwell time of 2.5 μs, and laser power of 58 μW (1PE) and 25 mW (2PE). Ca^2+^ traces are shown as changes in OGB-1 fluorescence (dF/F; excited at 488 nm, 1.7-5.2 μW). See **Figure S5B** for imaging/photostimulation schematics and for control trace without alloswitch. **C** Quantification of cell responses to agonist puffing from experiments as in panel **B. Top**, % of responding (colored) and silent (grey) cells during 2PE (red) or 1PE (violet) of alloswitch. Cells are shown as percentages of the total number 2 2+ of OGB-1 labeled cells detected in the imaged area (0.1 mm^2^). **Bottom**, The average peak amplitude of Ca^2+^ responses to 2PE in cells shown as responding in top graph (n = 69 cells, 3 animals) was about 15% lower (*p*<0.0001, unpaired t-test) than the amplitude of 1PE responses (n = 212 cells; 3 animals). Data are mean ± s.e.m.

In control experiments, photostimulation periods in naïve slices (no alloswitch, no agonist ejection; see **Figure S5C**) had no effect on cytosolic Ca^2+^. The efficacy of 2PE compared to 1PE of alloswitch in brain slices is in line with that observed in cultured cells (**Figure 3**), suggesting that diffusion of alloswitch isomers in brain tissue is not a limiting factor in photocontrolling mGlu_5_ receptor responses.

### Axial resolution to activate neurons and astrocytes in brain slices by 2PE of alloswitch is below 10 μm

Although the relevant parameter to evaluate the advantages of 2PE is the axial resolution achieved with a given compound, this value has not been determined for any of the previously reported azobenzene photoswitches, neither tethered^26,27^ nor freely diffusible^15^. In order to calculate the axial resolution of 2PE of alloswitch in brain slices, we recorded the responses of cells in one plane (OGB-1 fluorescence, z_img_) and compared the effect of photostimulation of alloswitch at five axial positions (z_stim_) within 15 μm above and below the imaging plane (**Figure 7A**). Additionally, the astrocytic marker sulforhodamine 101 (SR101, magenta in **Figure 7B**) was included in the ejection pipette to identify responding cells as astrocytes or neurons (only labeled with OGB-1, green in **Figure 7B**).

**Figure 7:**
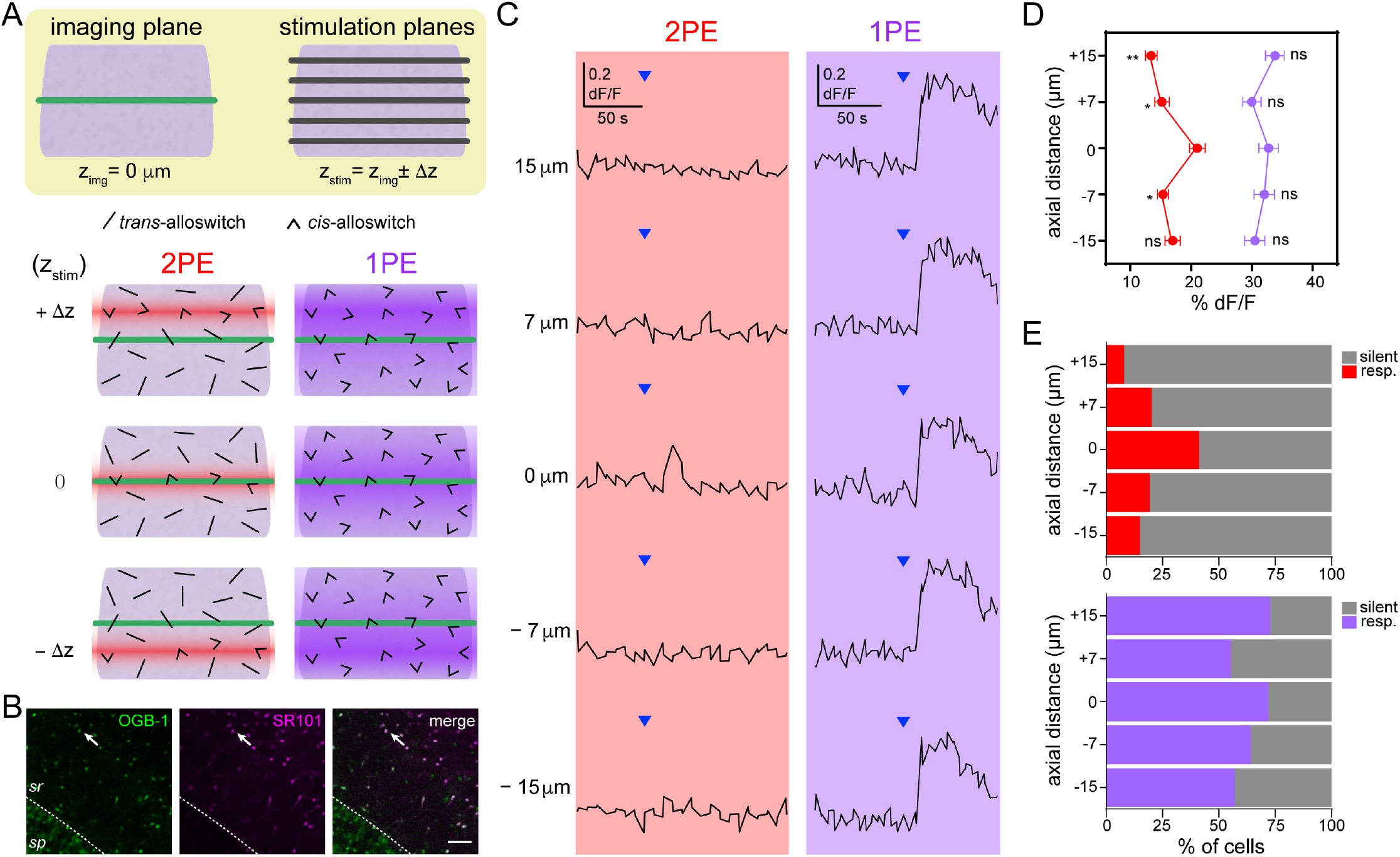
2PE of bath-applied alloswitch enables axially-resolved activation of endogenous mGlu_5_ in brain slices. **A** Experimental setup for imaging cells in the CA1 region of a rat acute brain slice at one plane (imaging plane, z_img_= 0 μm) while coincident 2PE (780 nm) and 1PE (405 nm) of alloswitch (10 μM) are performed at different positions inside the tissue along the z-axis (z_stim_= z_img_ ± Δz, with Δz= 0, 7, 15 μm) by moving the microscope objective. The 1P or 2P stimulation is done from the farthest z_stim_ (+15 or −15 μm) and sequentially approaching z_img_ in both directions. The position of the agonist ejection pipette is fixed throughout experiment. Photoswitching to *cis*-alloswitch using 2PE is restricted to a limited tissue volume compared to 1PE, so that 1PE photoactivates cells at z_img_ regardless of z_stim_, whereas 2PE only photoactivates imaged cells when Δz=0. **B** Fluorescent images of the pyramidal cell layer *(stratum pyramidale, sp)* and *stratum radiatum (sr)* of the hippocampus in a brain slice loaded with OGB-1 for Ca^2+^ imaging (left) and co-labeled with the astrocytic marker sulforhodamine 101 (SR101) included in the ejection pipette solution (middle). The merged image (right) allows distinguishing between astrocytes (OGB-1+/SR101+) and neurons (OGB-1+/SR101-) in Ca^2+^ activity movies. Arrows indicate one of the cells recorded, corresponding to traces shown in **C**. Scale bar is 50 μm. See quantification of cell types in **Figure S6**. **C** Representative Ca^2+^ responses from one cell of *stratum radiatum* (arrows in **B**). Agonist (blue arrowheads, 1 mM DHPG) was locally ejected and changes in OGB-1 fluorescence (% dF/F) at z_img_ were monitored at all indicated axial distances (z_stim_) for both 2PE (red box) and 1PE of alloswitch (violet box). See **Figure S7** for imaging/photostimulation schematics and for control injections with/without alloswitch and before/after photostimulations. **D** Average peak amplitudes of Ca^2+^ responses obtained from z_img_ while photostimulating cells at the indicated axial distances (Z_stim_ = +15, +7, 0, −7 and −15 μm). Peak amplitudes of responses acquired at z_img_ while photostimulating at z_stim_= ±7, ±15 were compared to that of z_stim_=0. 2PE of alloswitch along the Z axis evoked the largest response when z_stim_= 0 (in % dF/F, 13.5 ± 1.0, 15.2 ± 1.2, 21.0 ± 1.3, 15.4 ± 0.9 and 17. 0 ± 1.3, respectively). 1PE responses did not significantly change with Z_stim_ (in % dF/F, 33.8 ± 1.6, 30.0 ± 1.5, 32.7 ± 1.6, 32.0 ± 1.7 and 30.5 ± 1.7). Data are mean ± s.e.m; two-way ANOVA, \**p*<0.05, \*\**p*<0.01; ns, not significant. A full-width at half-maximum (FWHM) of 6.2 ± 2.2 μm for 2PE was obtained by fitting the response amplitude during 2PE to a Gaussian function, and gives the approximate resolution of 2PE of alloswitch that can be achieved under these experimental conditions. **E** Percentage of silent (grey) and responding cells (colored) observed at z_img_ during 2PE and 1PE of alloswitch at each indicated z_stim_. The number of responding cells within the imaged area (0.1 mm^2^) during 2PE was 9, 23, 47, 22 and 17 (**top**; in %, 7.9, 20.2, 41.2, 19.3 and 14.9) and during 1PE, 83, 63, 82, 73 and 65 (**bottom**; in %, 72.8, 55.3, 71.9, 64.0 and 57.0). A full-width at half-maximum (FWHM) of 10.3 ± 2.3 μm for 2PE was obtained by fitting the percentage of cells responding during 2PE to a Gaussian function.

We first quantified photoresponses of all neurons and astrocytes. 1PE produced Ca^2+^ responses in the imaging plane regardless of the position of photostimulation plane (**Figure 7C right**), in agreement with the results observed in cell culture (**Figure 4**). By contrast, 2PE responses were observed mostly when photostimulation was performed on the imaging plane (‘0 μm’ label in **Figure 7C left**), indicating that mGlu_5_ receptors silenced with *trans*-alloswitch (see **Figure S7**) were selectively rescued by photoisomerizing alloswitch to the *cis* form in a confined region in the axial direction. The Ca^2+^ responses to 1gPE and 2PE were quantified as a function of the axial distance in two ways. First, the response amplitude expressed as relative change in fluorescence (%dF/F) was around 30% and did not change significantly with the distance for 1PE, whereas it showed a maximum of 20% at 0 μm for 2PE (**Figure 7D**). Fitting 2PE data to a Gaussian function yielded a full-width half maximum (FWHM, axial resolution; see **Methods 6.4** for details) of 5.4 ± 0 μm. Second, the percentage of cells displaying photoresponses was approximately 60% at any axial distance for 1PE, whereas 2PE responses were maximal at 0 μm (40% of cells, **Figure 7E**) and the FWHM of this Gaussian distribution was 10.3 ± 2.3 μm.

Co-labeling with SR101 indicated that about half of the cells (52%) responding to 1PE of alloswitch were astrocytes (**Figure S6B–D**), supporting the use of alloswitch as a versatile tool to photocontrol different cell types in intact brain tissue. Cells responding to 2PE included a higher fraction of astrocytes (63%) but no differences between astrocytes and neurons were observed in the amplitude or time course of 1PE or 2PE responses in our experimental conditions (see example traces in **Figure S6C**).

## Discussion

Existing strategies to elucidate protein function include genetic or pharmacological manipulations, however both show several limitations. In first place, genetic targeting is usually aimed at either ablate protein expression via antisense oligonucleotides, or disrupt the gene encoding for the protein under study, either in the whole organism in knock-outs or at specific ages, tissues and cell types in conditional knock-outs. All experimental observations based on these approaches rely on the indirect effects derived from receptor absence, which are wider and less specific than the direct effects of receptor activation/inhibition. Recent advances in these fields have raised concerns about the mechanisms enabled by genetically modified organisms to compensate for or adapt to protein loss^41^, and how such compensatory mechanisms influence the outcomes of experimental observations is largely underestimated^42^.

In second place, pharmacological blockade of protein function is a fast, simple and flexible methodology that is routinely used to manipulate receptor activity without altering genes or protein levels. Although continued efforts in drug development are filling the gaps in the availability of potent and receptor-selective ligands, pharmacological agents are still limited by the relatively slow time course of compound application, diffusion, and removal, reducing their experimental versatility^3^. Strategies to constrain the spatial and temporal functions of drugs include localized perfusion and fast delivery systems, and conjugation with macromolecules^43^. However, these methods fall short of the spatiotemporal scales relevant to neural function, such as the micrometric dimensions of dendritic spines and astrocytic processes, neuronal action potentials and astrocytic Ca^2+^ signals lasting milliseconds to seconds. In contrast, photopharmacology allows control of drug activity at the spatiotemporal coordinates of the patterned illumination defined by the user. It encompasses a new class of diffusible ligands that are light-regulated and enable turning on and off the activity of endogenous receptors at physiological locations and time-scales.

Advanced imaging methods using patterned illumination by multiphoton microscopy with femtosecond pulsed lasers^23^ enabled imaging the cortical activity in mice at millimeter depths^44^. Recent progress in 2PE techniques including scanless spatial light modulation and holographic patterning^45,46^, as well as temporal focusing techniques^47^ are currently being exploited for optogenetic stimulation^48–52^. However, applications to photopharmacology have been reported only in a few cases, and focused on molecular aspects rather than functional outcomes^15,26,27^.

To take advantage of the spatiotemporal control of light regulated drugs using 2PE (‘two-photon pharmacology’, 2PP) we chose alloswitch, whose nanomolar activity and subtype selectivity make it especially suitable to manipulate endogenous receptors in intact neural tissue. Bath application of alloswitch immediately silenced mGlu_5_ receptors, and 2PE produced rapid and localized rescue of mGlu_5_ agonist responses. Illumination power, wavelength and other 2PE conditions can be achieved with commercial pulsed lasers (**Figure 2**) and were effective in all tested alloswitch analogs (four selected from a library of 18,^28^; **Figure S3**). 2PE of alloswitch achieved mGlu_5_ responses comparable to endogenous activity induced by agonist, yet weaker than 1PE in the same conditions (**Figure 3**). Nevertheless, screening the 2PE responses of available photopharmacological analogs^28^ and designing new compounds with improved 2PE isomerization efficiency (Cabré, Garrido-Charles *et al.,* in review) should give rise to a new generation of 2PP-optimized ligands able to transduce complex three-dimensional illumination patterns into selective activation of endogenous receptors in neuronal circuits with total reliability. Compared to caged ligands^53^, alloswitch and other reversible photoswitchable compounds provide better receptor subtype selectivity and allow controlling the receptor deactivation rate^4^.

The efficiency of 2PE of alloswitch observed in our functional study using raster scanning (**Figure 3D**) suggests that the absorption and quantum yield of the molecule in the 2PE regime is higher than that of other azobenzene-based photoswitches^15,26,27^. This effect might be due to the favorable push-pull character of alloswitch molecules, but downstream amplification of the signal by G protein-coupled receptor (GPCR) effector proteins cannot be ruled out^27^. The proportion of cells in the field-of-view responding to 2PE compared to 1PE in brain slices (33%, **Figure 6C top**) is in line with that observed in cultured cells (36% at 0 μm, **Figure 4D**). This suggests that diffusion of alloswitch isomers in the highly-packed brain tissue^54^ is not a limiting factor to photocontrol mGlu_5_ receptor responses, and that improving the 2PE molecular properties of alloswitch should have a direct effect on 2PP performance. Overall, the results of **Figures 5–6** constitute the first demonstration that the activity of neuronal receptors can be controlled by 2PE of allosteric drugs in intact brain slices. Interestingly, using 1PE of alloswitch, Ca^2+^ responses were recovered in a significantly higher fraction of cells in the field of view than those responding to DHPG in the absence of alloswitch (**Figure 5E**). Similar effects are observed in cell lines (**Figure 3D**) and have been previously reported^16,28^. In brain slices, these results suggest that pharmacological silencing of mGlu_5_ receptors followed by rapid photo-rescue in a physiological setting uncovers neglected features of receptor balancing in response to prolonged arrested signaling. Dedicated studies are required to investigate the relevance of this phenomenon.

The quantification of mGlu_5_ responses yielded an axial resolution for 2PE of alloswitch in brain slices between 5 and 10 μm (either quantifying the response %dF/F or the number of responding cells in **Figures 7D and 7E**, respectively).

These values match well the average size of brain cell somata, and are comparable to those obtained using optogenetic tools (12.5 μm axial resolution with C1V1 channelrhodopsin;^55^). In our wide-field, low-magnification fluorescence microscopy movies, fluorescence changes are observed mainly at the cell bodies (Movies S3–4). Although the presence of TTX in the bath guaranteed that the Ca^2+^ activity was initiated by resident mGlu_5_ receptors and not propagated, the resolution obtained with 2PE is still influenced by the spatial distribution of the cells expressing mGlu_5_ in brain slices, and by the limited sensitivity of chemical dyes to measure Ca^2+^ responses in cellular processes away from the soma. Imaging and photostimulation at higher magnifications than those used in the present study leave room to improve the spatial resolution of 2PP. For example, Ca^2+^ in distal dendritic branches or peripheral astrocytic processes could be efficiently monitored using genetically-engineered Ca^2+^ indicators (GECI), and mGlu_5_ targeted using alloswitch molecules by first silencing receptors widely, and then selectively rescuing them by 2PE in micrometric volumes corresponding to single spines or individual astrocyte processes.

Alloswitch photoresponses observed in astrocytes and neurons (**Figures 7B and S6**) are in agreement with the reported expression of functional mGlu_5_ in both cell types^39^. The slightly lower fraction of neurons (compared to astrocytes) that respond to 2PE compared to 1PE may be due their size and/or spread of astrocytic branches being larger than the volume effectively illuminated by 2PE, and is in accordance with the high complexity of Ca^2+^ signals displayed by astrocytic compartments^56^. Further insights into mGlu_5_ signaling in neurons and astrocytes could be drawn from higher magnification studies of brain activity with 2PP. For example, our raster-scanned, plane-selective activation of glial and neuronal mGlu_5_ receptors in intact neural tissue could be enhanced using 2PE holography to achieve full threedimensional patterned control at the single cell or subcellular scales, and provide an unprecedented view over the spatial spread and time course of endogenous receptor signaling^57,58^.

The robustness of mGlu_5_ silencing and 2PE-rescue of responses by alloswitch is outstanding both in cultured cells and in brain slices (**Figures 4 and 6**). Given the nanomolar potency of *trans*-alloswitch, its ability to completely silence mGlu_5_ activity was not a matter of concern in any scenario. However, that particular property together with the expected partial efficacy of 2PE to convert *trans*-alloswitch to the *cis* form suggested that a large fraction of *trans*-alloswitch would remain in the focal volume, and that it would be surrounded by an even larger pool of *trans* isomers ready to flood that volume by diffusion. In this context, the fact that mGlu_5_-specific intracellular signaling can be elicited in the presence of alloswitch during periods of 2PE demonstrates that *trans* and *cis* alloswitch do not interfere with one another in these experiments. This could be due in part to limitations in the diffusion of *trans*-alloswitch into the 2PE volume. However, this explanation is unlikely because 2PE of alloswitch was also observed in cultured cells, where diffusion of molecules is not hindered by anatomical constraints as in neural tissue^54^. Another possibility is that mGlu_5_ receptor-bound *trans* molecules could be photoconverted to *cis* inside the allosteric binding pocket of the protein, and if photoconversion (picoseconds) was faster than the kinetics of unbinding from the receptor (milliseconds or more; see^59,60^), then the 1 nm-wide entrance of the allosteric ligand binding site of mGlu_5_^61^ might hinder the exit of *cis* molecules and prevent *trans* molecules from entering and inhibiting the receptor. This hypothesis could be tested using FRET measurements of receptor conformational changes^62^ combined with methods for rapid photoswitching^4^.

Caged compounds are another class of photopharmacological agents that can be activated using 2PE. Because irreversibly photoreleased molecules cannot be deactivated, their effect can only cease by diffusion in the surrounding medium (which is maximal when the 2PE volume is small) or by neurotransmitter removal via enzymatic hydrolysis and/or reuptake inside cells. In contrast, reversible photoswitches like alloswitch are in steady-state equilibrium between their active and inactive isomers. This equilibrium can be displaced in either direction by means of illumination or thermal relaxation. As the diffusion of molecules is constrained at the nanoscale in the synaptic cleft^54^, reversibility constitutes an important advantage of photoswitching compared to irreversible uncaging of ligands.

Signaling pathways involving mGlu_5_ function govern important physiological processes whose mechanisms remain controversial due to the lack of suitable experimental tools to study them. The pharmacologically selective and spatiotemporally controlled activation of mGlu_5_ in intact brain tissue that is demonstrated here will be useful to address several biological questions. For example, in the developing cortex, is glutamatergic signaling inducing specific spatiotemporal patterns of mGlu_5_ activation that organize neurons and glia into functional networks^63,64^? After development, what patterns of mGlu_5_ activation drive synaptic plasticity^65^, homeostatic scaling^66^, and regulation of basal transmission^67^? In addition, what are the roles played by the different mGlu_5_-expressing cell types of the hippocampus in the mechanism of mGluR-dependent long-term depression^68^? How many spines, dendritic branches, or cells are involved in synaptic plasticity^69^? Further increasing the resolution of photostimulation will allow studying how mGlu_5_ receptors expressed on organelles (including nuclear membranes and other subcellular localizations;^70^) contribute to the complexity of canonical and non-canonical signaling pathways activated downstream of mGlu_5_^71^.

Thus, 2PP in physiologically relevant settings such as brain slices, or in anaesthetized and awake animals will be an invaluable tool to understand how neurotransmitter receptors of neurons and glia function in the intact brain.

### Conclusions

We have demonstrated that photoswitchable allosteric modulators of mGlu_5_ receptor activity (alloswitch and its analogs) can be effectively photoisomerized using pulsed near-infrared lasers (2PE) in cultured cells and acute brain slices. Combining potent and subtype-selective photopharmacology with 2PE allows us to silence and photo-rescue endogenous receptors rapidly and reversibly. We calculate the axial plane selectivity of 2PE of a photoswitchable ligand for the first time in brain slices (10 μm), and this value corresponds to the size of single neuronal and astrocytic somata. Thus, photoswitchable ligands of endogenous receptors allow high-resolution two-photon pharmacology (2PP), despite diffusion in neural tissue. This method does not require any genetic manipulation, and is readily applicable to intact tissue. Its efficiency to induce mGlu_5_ photoresponses indicates that the absorption and quantum yield of alloswitch in the 2PE regime is higher than that of other azobenzene-based photoswitches. All alloswitch analogs tested in this study were found to be 2PE-active, which should allow further optimization of their 2PE properties. In addition, the activation of neuronal and glial mGlu_5_ receptors can be traced separately in intact tissue, which opens the way to study their distinct physiological roles with unprecedented spatiotemporal resolution and tissue depth. In this way, 2PP is a useful and versatile addition to the molecular toolbox to study neural function.

## Methods

### 1 Diffusible photoswitches and other drugs

Freely-diffusible photoswitchable allosteric modulators of mGlu_5_ receptors were obtained as previously described for alloswitch^16^ and compounds **1c-d, 1j** and **6**^28^. Other drugs used (quisqualate, DHPG, LY367385, TTX) were purchased from Tocris.

### 2 Microscopy, light sources, and general imaging conditions

Imaging of cultured cells and brain slices, and photostimulation of alloswitch and its analogs were performed using an inverted laser-scanning confocal microscope (TCS SP5, Leica Microsystems) equipped with a HCX PL APO 40×/1.25-0.75-NA oil objective for imaging cultured cells, and a HC PL APO 20×/.0.7-NA CS air objective (Leica Microsystems) for imaging acute slices. The pulsed laser (PL) used for two-photon excitation (2PE) of alloswitch and Ca^2+^-imaging was an IR laser MaiTai Wide Band (710nm–990nm, Spectra Physics). Continuous-wave lasers (CWLs) used were: a LASOS 405-50 50 mW 405-nm laser diode for one-photon excitation (1PE) of compounds, and a LASOS LGK7872 65 mW Argon laser for visualization of OGB-1, GCaMP6s, or eYFP-tag on cells (488/514/561-nm lines, respectively). For all wavelengths, images were acquired by bidirectional raster scan at 400 Hz. For cultured cells, image size was 387.5×387.5 μm, pixel size 1.52 μm, and pixel dwell time 9.8 μs. For slices, image size was 775×775 μm and 512×512 pixels; pixel size and dwell time were 1.52 μm and 4.9 μs (or 3.04 μm and 9.8 μs for 256×256 pixels images). For experiments where a 2.5× optical zoom was applied, image and pixel size were 310 μm and 606.65 nm respectively, whereas dwell time was unchanged. Emitted fluorescence was collected using hybrid detectors. Frame rates and laser powers used for specific experiments are detailed in the corresponding sections.

### 3 Cell culture procedures and imaging

#### 3.1 Plasmids

For “all-2P” experiments (2P-imaging of Fura-2 combined with 2P-photostimulation of alloswitches), we used a plasmid where the mGlu_5_ receptor was in fusion with a fluorescent reporter (mGlu_5_-eYFP;^72^) to visualize expressing cells and identify false-positives for photostimulation. For experiments combining 1P-imaging with 1P- or 2P-photostimulation, we co-transfected cells with a plasmid for GCaMP6s under a ubiquitous promoter for Ca^2+^-imaging (Addgene, plasmid #40753) and a plasmid encoding for the mGlu_5_ receptor lacking eYFP^16^ to avoid bleed-through into the Ca^2+^-reporter channel.

#### 3.2 Cell culture and transfection

HEK tsA201 cells were maintained at 37 °C and 5% CO_2_ humidified atmosphere in Dulbecco’s Modified Eagle Medium/Nutrient Mixture F-12 (Gibco DMEM/F-12, Thermo Fisher Scientific 11320) with 10% Fetal Bovine Serum (Gibco heat-inactivated FBS, Thermo Fisher Scientific 10500-056-100ml) and 1% antibiotics (Penicillin/Streptomycin, Sigma-Aldrich P4333). We induced transient expression of the plasmids in 10^6^ cells resuspended in 2 ml using the polymer X-tremeGENE 9 DNA Transfection Reagent (Roche Applied Sciences 06365787001) following manufacturer’s instructions (1 μg total DNA and 3 μl reagent in 100 μl FBS-free culture medium). The Ca^2+^ reporter was co-transfected with the receptor in a proportion of 1:1, with other conditions unchanged. Cells were seeded onto 21-mm glass coverslips treated with Poly-L-Lysine (Sigma-Aldrich P4832) to allow cell adhesion. Pre-confluent cultures were used for experiments at 48 to 96 hours after transfection. Coverslips were mounted on an Attofluor cell chamber (Thermo Fisher scientific A7816) in bath solution containing (in mM): 140 NaCl, 5.4 KCl, 1 MgCl_2_, 10 HEPES, 10 D-glucose and 2 CaCl_2_; pH was adjusted to 7.40 using NaOH.

#### 3.3 “All-2P” experiments workflow (2P-imaging of Fura-2 in mGlu_5_-eYFP cells)

##### 3.3.1 Fura-2 loading

Cells expressing mGlu_5_·eYFP were loaded with the Ca^2+^-indicator Fura-2 AM (10 μM, Thermo Fisher Scientific F1201) diluted in culture medium and 1% DMSO for 30 minutes in the cell incubator (37 °C and 5% CO_2_), and then rinsed.

##### 3.3.2 Identification of mGlu_5_-eYFP-expressing HEK cells

To double-check for mGlu_5_+ cells, fluorescence images of the eYFP-tag on the receptor were acquired using the 514-nm line of the Argon laser or the pulsed laser set at 900 nm.

##### 3.3.3 2P Ca^2+^-imaging using Fura-2

Fura-2 was excited with the pulsed laser at a frame rate of 0.2 Hz at NIR wavelengths (760, 780, 800, or 820 nm, see **Figure S1** for example traces), approximately corresponding to twice the wavelength (380–410 nm) of 1P excitation with continuous-wave lasers. Fura-2 was used in the single-wavelength (non-ratiometric) modality, to accommodate fast imaging rates of the Ca^2+^ indicator. Using this setup, increases in intracellular Ca^2+^ are detected as decreases in the fluorescence emission of Fura-2 (consistent with its emission spectra following 380 nm excitation; see **Figure S1A**). Laser power was set to the minimal value allowing satisfactory visualization of Fura-2 (~3 mW on sample for all wavelengths used), to minimize potential 2PE-isomerization of alloswitch during imaging. Emitted fluorescence was collected using hybrid detectors with spectral range 420–515 nm and standard gain settings.

#### 3.4 1P-imaging of GCaMP6s/mGlu_5_ co-transfected cells

GCaMP6s fluorescence was monitored using the 488 nm line of the Argon laser, with light intensity adjusted to the lowest power allowed by the system (100 nW on sample) to avoid *cis-to-trans* photoisomerization of alloswitch. Images of GCaMP6s were acquired at a frame rate of 0.2 Hz. The pinhole of the confocal system was set at the maximum aperture (optical section thickness was 7.4 μm), and the emitted fluorescence was collected using hybrid detectors (490-550 nm). In a set of experiments, cell morphology and thickness was detected by imaging GCaMP6s at 900 nm using the pulsed laser.

### 4 Acute rodent brain slices procedures and imaging

#### 4.1 Ethical statement

Procedures were performed in accordance with the European guidelines for animal care and use in research, and were approved by the Animal Experimentation Ethics Committee at the University of Barcelona (OB-432/16).

#### 4.2 Dissection of acute brain slices

Acute brain slices were obtained from Sprague-Dawley rats (Envigo) of 6 to 15 postnatal days (P6–15). Animals were sacrificed by decapitation to remove the brain. Coronal slices (350-μm thick) were obtained in ice-cold cutting solution containing (in mM) 27 NaHCO_3_, 1.5 NaH_2_PO_4_, 2.6 KCl, 222 sucrose, 2 MgSO_4_ and 2 CaCl_2_ and saturated with 95% O_2_ and 5% CO_2_, using a vibrating-knife microtome (1000S Vibratome, Leica). Hippocampal slices were collected in a holding chamber filled with artificial cerebrospinal fluid (aCSF) containing (in mM) 126 NaCl, 26 NaHCO_3_, 1.145 NaH_2_PO_4_, 3 KCl, 10 glucose, 2 MgSO_4_ and 2 CaCl_2_, saturated with 95% O_2_ and 5% CO_2_. Slices were recovered in the holding chamber for at least 1 hour at RT before OGB-loading, and used not more than 7 hours after sacrifice.

#### 4.3 Ca^2+^-imaging of acute brain slices using OGB-1

##### 4.3.1 Bulk-loading of the Ca^2+^-indicator Oregon Green BAPTA-1 AM

A group of 4 slices was loaded with 20 μM OGB-1 (Life Technologies O6807) in oxygenated aCSF containing 0.04% (w/v) Pluronic F-127 (Sigma P2443) and 0.5% DMSO (Sigma 276855). Slices were loaded during 15 minutes at 30°, and for 10 more minutes at RT, and then returned to the holding chamber.

##### 4.3.2 1P Ca^2+^-imaging in slices

For imaging, a slice was transferred into a 35-mm dish chamber (MatTek, P35G-1.0-14-C) along with a dish insert (Warner Instruments, RC-33DL) to allow perfusion with oxygenated aCSF (see ***Methods 4.2*** for aCSF composition) complemented with 100 μM LY367385 (mGlu_1_ antagonist, Tocris 1237) and 1 μM TTX (Na_V_ channel blocker, Tocris 1069) to isolate Ca^2+^-events solely induced by mGlu_5_ activation (hereafter: *aCSF+LY/TTX),* and recirculated at 1.5-2 ml/min (Reglo peristaltic pump, Harvard Apparatus, 73-2449) at RT. Calcium events were recorded in *stratum oriens* to *stratum lacunosum-moleculare* of the CA1 region of the hippocampus by exciting OGB-1 with the 488 nm Argon laser set at a frame rate of 1–0.33 Hz and low power (1.7–5.2 μW) to avoid unwanted *cis-to-trans* isomerization of alloswitch. The confocal pinhole was set to obtain optical sections of 6.182–17.5 μm.

##### 4.3.3 Pressure-ejection of the mGlu_5_ receptor agonist in slices

Micropipettes with a tip resistance of 3–6 MΩ were prepared from 1.2/0.69-mm OD/ID borosilicate glass capillaries (World Precision Instruments, GC-120F-10) using a pipette puller (Sutter Instruments, P-97), and filled with *aCSF+LY/TTX* (detailed in ***Methods 4.3.2***) containing 1 mM (S)-3,5-DHPG, or 2 mM (R,S)-3,5-DHPG (mGlu_1/5_ agonist, Tocris 0805 and 0342 respectively), and the astrocyte marker Sulforhodamine 101 (40 μM, SR101, Sigma S7635). Pipettes were held with a micromanipulator (Narishige MN-153) off the microscope stage, and placed in contact with the slice surface on top of the field of view in a shadow-guided manner using transmitted light. Agonist ejection was initiated manually using a pressure-ejection system (Eppendorf FemtoJet, VWR) set at 500–800 hPa during 1 s.

##### 4.3.4 Identification of SR101+-cells

SR101^+^-cells were visualized at 561 nm with the Argon laser to confirm ejection site and identify cell types.

### 5 Photostimulation of diffusible photoswitches

#### 5.1 Bath-application of compounds

In cultured cells, agonist (3 μM quisqualate, Sigma-Aldrich Q2128) and photoswitches (1 μM from 1 mM stocks in DMSO; total DMSO was 0.1%) were supplemented as described previously^16^. In brief, compounds were added manually during image acquisition in the recording chamber by thoroughly pipetting a small volume to assure a good mixing of the solutions. In slices, recirculating *aCSF+LY/TTX* was replaced with oxygenated *aCSF+LY/TTX* containing 10 μM alloswitch (in 0.02% DMSO) to block responses of cells to puffing of the mGlu_5_ agonist. Alloswitches were incubated for at least 7 minutes in both culture and slice experiments to allow silencing of mGlu_5_ receptors before photostimulation.

#### 5.2 Photostimulation

Photostimulation at 1PE or 2PE was achieved by raster-scanning the field-of-view with the corresponding lasers, executed through the same light path used for Ca^2+^-imaging. To do this, raster-scans for photostimulation were interleaved between two consecutive image acquisitions (= imaging cycle) of either Fura-2, GCaMP6s, or OGB-1.

Photostimulation scans were repeated consecutively 13 and 10 times per imaging cycle in Fura-2-loaded and GCaMP6s-expressing cells, respectively, and 1 to 3 times per imaging cycle in slices. Image and pixel size, and pixel dwell time for photostimulation were the same as for imaging (see section **Methods 2** and diagrams in **Figures S1, S4-5 and S7**). Adjustments to the frame rate were introduced by the imaging software to accommodate for hardware changes occurring during video recordings *(i.e.* switching light wavelength and power, and stage or objective movements), and accounted for imaging frame rates reduced to ~0.15 Hz during 2PE in Fura-2-loaded cells, ~0.17 (1PE) and ~0.12 Hz (2PE) in GCaMP6s-cells, and average values of 0.3–0.4 Hz (1PE) and 0.3 Hz (2PE) in slices. All adjustments were recorded and considered at the time of data analysis and plotting, but are not represented in supporting Movies.

##### 5.2.1 1PE

One-photon excitation (1PE) was accomplished using the 405 nm diode laser set at 2 μW in GCaMP6s-HEK cells, and 58-98 μW in slices (power measured after objective lens).

##### 5.2.2 2PE

The light source for two-photon excitation (2PE) was the pulsed IR laser MaiTai Wide Band tuned at 780 nm, unless otherwise indicated. Laser power was adjusted to be 12 mW on sample in experiments with Fura-2-loaded cells, 20 mW for GCaMP6s-expressing cells, and 25 mW for slices (powers measured after objective lens using a thermal-head power sensor; L30A-V1, Ophir Photonics). Note that in experiments where 2PE of compounds was combined with 2P-imaging of Fura-2, the software used for imaging and photostimulation arbitrarily increased the light intensity set for the imaging scans, which resulted in increased fluorescence of the sample, but did not correspond to a real change in intracellular Ca^2+^, and neither masked the photostimulated Ca^2+^-oscillations (as illustrated in **Figure S1D-E**). This increase in baseline fluorescence was not *post-hoc* corrected in any of the cell traces shown.

##### 5.2.3 Photostimulation at axial distances other than the focal plane

In experiments with cultured cells, photostimulation at varying axial distances was achieved by moving the motorized stage downwards, thus shifting the focal plane for 1PE or 2PE above the cells and inside the bath containing the photoswitches. For slice experiments, the presence of the ejection pipette prohibited any stage movement, thus photostimulation at axial distances above and below the imaging plane was achieved by moving the objective in either direction. Image settings and light powers were as already detailed above.

### 6 Image analysis and statistics

Acquired image sequences were stored in the Leica image format and used to extract exact times of image acquisitions in comma-separated values (.csv), and image stacks for offline analysis with ImageJ software (NIH release: Nov 2013).

#### 6.1 Analysis of Fura-2 fluorescence and 2PE in cultured cells

Cell profiles were defined in ImageJ, and the average fluorescence intensity for each time-point was extracted (F). For each identified cell, fluorescence changes (indicated as dF/F in the figures) relative to the first frame of each series (F_0_) were calculated as (F-F_0_)/F_0_ and plotted in Excel with a custom-written VBA macro. Cells showing Ca^2+^-events after bath-application of alloswitches and before photostimulation, if any, were excluded from further analysis. Cells displaying a single Ca^2+^-event during 2PE were not considered as successfully excited. Only cells showing two or more Ca^2+^-peaks were classified as responsive and considered for counting the number of peaks (# oscillations), identifying peak time of first, second and last oscillation observed (first time-point where the fluorescence signal clearly decreased relative to basal, indicating an increase in the intracellular Ca^2+^, and before the minimum fluorescence was observed, corresponding to the peak amplitude of the Ca^2+^ response). These values were used to calculate the delay for the appearance of the first peak from the photostimulation onset (delay), the time between first and last peak (duration) and their average frequency (= # oscillations/duration). After all responding cells had been assigned a delay time and oscillatory frequency, we loaded the eYFP channel images to cross-check for transfected cells. No responding cells were found to lack mGlu_5_-eYFP expression.

#### 6.2 Analysis of GCaMP6s-fluorecence and 1PE/2PE in cultured cells

Cell profile and fluorescence intensities were obtained as described in the previous section (**Methods 6.1**) for all cells displaying fluorescence changes at any time-point. Cells with unstable fluorescence in the absence of stimulation were discarded. In experiments where both 1PE and 2PE were done sequentially on the same cell, cells were further sorted if showing inconsistent oscillatory responses to 1PE, defined as either having ≤2 peaks/5 minutes, or when 1PE responses were not successfully reproduced in the second 1PE photostimulation (1PE at the focal plane). The remaining cells were classified as *‘1P-oscillating’* or *`1P-CaEvents`* and pooled to count total number of cells analyzed. Cells were further classified in *‘2P-oscillating-0μm’*, ‘*2P-CaEvents-0μm`*, ‘*2P-CaEvents-10μm’* or *‘2P-NOresponse’*, to determine % of photoactivated (‘*oscillating*’ + *‘CaEvents’)* and oscillating cells (‘*oscillating*’ only) over the total number of cells considered for analysis. No cells were found active at 2PE and inactive at 1PE. No cells were found to display Ca^2+^-events when 2PE was done 30 μm above the cell plane, or events in the form of oscillations 10 or 30 μm away from the cells, and the corresponding categories were not created. For both experiment types, traces of cells showing oscillatory behaviors were analyzed using Igor Pro (WaveMetrics) to determine: first time-point above basal Ca^2+^ (t_ON_), number of oscillatory events observed (# oscillations), and last time-point before return to baseline or stop of oscillatory behavior (t_OFF_). These data were used to calculate the time delay between the beginning of the photostimulation and the onset of the oscillatory behavior (delay), overall duration of the oscillatory behavior (duration = t_ON_ – t_OFF_), and number (# oscillations) and average frequency of calcium oscillations (freq. of oscillations = # oscillations / duration). Graphs and statistics were generated with Prism (GraphPad).

#### 6.3 Analysis of OGB-1 fluorescence and 1PE/2PE in slices

To assist OGB fluorescence image analysis for 2PE experiments in slices, an ImageJ macro was written at ADMCF (Advanced Digital Microscopy Core Facility, IRBB) which detects cell somata after noise reduction and registration (as required), extracts fluorescence changes relative to baseline (dF/F) over time, and classifies cells as responsive versus non-responsive based on the presence or absence of significant fluorescence peaks above noise following stimulation. Code and detection pipeline, together with a sample time-lapse are available online (https://sites.google.com/a/irbbarcelona.org/adm/image-j-fiji#TOC-Somata-segmentation-and-time-response-analysis-in-acute-rodent-brain-slices-). For all detected regions-of-interest (ROIs), exported dF/F traces were graphed using a custom-written macro in Excel. Responding and non-responding cells were validated manually for each condition, and then counted as percentages relative to the total number of cells detected in the field of view. First, ROIs displaying spontaneous activity (fluorescence spikes before stimulation) or poor signal-to-noise ratios were manually excluded from further analysis; second, when detectable changes in fluorescence where observed over 20 to 30 frames after agonist ejection, cells were counted as “responding”, and otherwise classified as “silent”; finally, the peak amplitude of responding cells was extracted as the maximum dF/F value observed within this frame range. Detection of ROIs with ImageJ, dF/F calculation relative to first frame fluorescence, and cell classification were done manually for a set of lower magnification 1PE experiments (**Figure 5**) using the same criteria. For experiments where efficacy (number of responding cells) and efficiency (peak amplitude of the cell response) of photoactivation at the focal plane were measured along with those of photostimulation at 4 other axial distances (**Figures 7** and **S7C**), cells that were insensitive to either DHPG or alloswitch-1 (**Figure S7B**) were excluded from further analysis.

#### 6.4 Data and statistics

Data are represented as mean ± s.e.m. unless otherwise indicated. Number of cells, slices and animals used are indicated in figure captions. Graphs were generated with Igor Pro 6 (WaveMetrics) or Prism 5 (GraphPad) and figures with Illustrator CC (Adobe). Statistics performed using Prism software are indicated in figure captions. Gaussian fittings were done using Origin software using data points with amplitudes significantly different from the peak. The full width at half maximum (FWHM) was calculated by multiplying the standard deviation of the fitted curve by a factor of 2.355.

## Acknowledgements

This research has received funding from the European Union’s Horizon 2020 Framework Programme for Research and Innovation under the Specific Grant Agreement No. 720270 (Human Brain Project SGA1), and the ERA-Net programme (Modulightor project). We also acknowledge financial support from AGAUR/Generalitat de Catalunya (FI fellowship, CERCA Programme and 2017-SGR-1442 project), FEDER funds, MINECO/FEDER (project CTQ2016-80066-R), Fundaluce foundation, and Ramón Areces foundation. SP was supported by a FI fellowship (2014FI_B2 00160). HL was supported by a fellowship of the IBEC International PhD Programme. MB was supported by a Marie Curie Reintegration Grant (H2020-MSCA-IF). KEP receives support from NIH/NINDS (R01NS099254) and NSF Biophotonics (1604544). We thank Jordi Hernando (Autonomous University of Barcelona) for useful discussions on 2PE, Pere Català (Utrecht University) for help with GCaMP experiments, Francisco Ciruela (University of Barcelona) for sharing the mGlu_5_-eYFP plasmid, Erin Schuman (Max-Planck Institute for Brain Research, Frankfurt) for supporting preliminary 2PE experiments, and Ashraf Muhaisen (Institute of Neuroscience, University of Barcelona) for support with slice preparation.

## Author contributions

SP, GP and PG designed the experiments. SP and HL conducted all experiments and analyzed data. SP, A. Lladó, LB and JC set up the local perfusion system and inverted confocal microscope for brain slice imaging and stimulation. GP, MB and KP conducted preliminary experiments in slices. ST wrote the ImageJ scripts to analyze neuronal activity. XGS and A. Llebaria contributed reagents. ES contributed equipment for slice experiments. SP and PG wrote the original draft. All authors reviewed and edited the paper. PG conceived and supervised the project.

## Supplemental Information

### Supplemental Figures

**Figure S1 (related to Figure 1):**
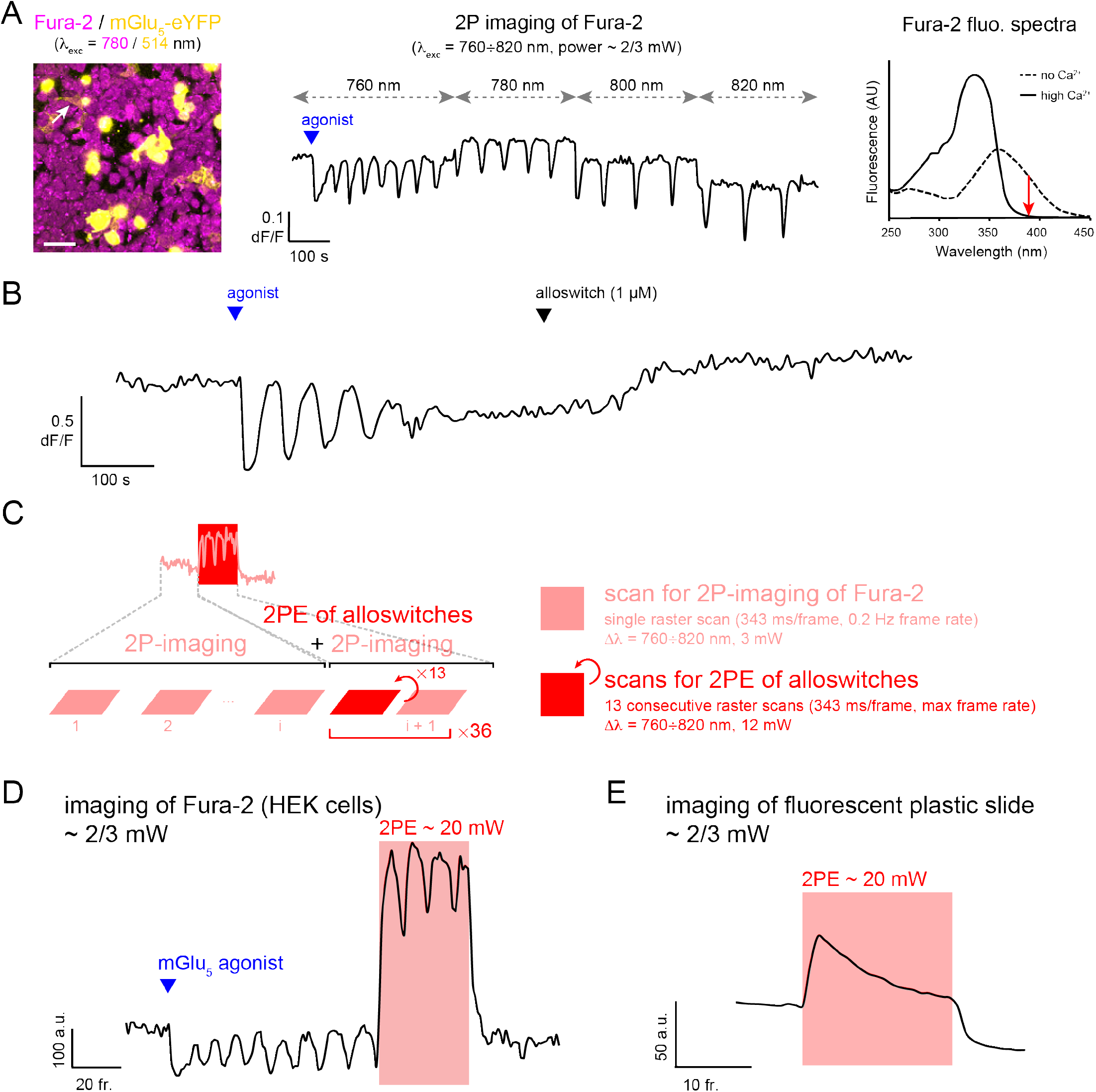
Panel A: Single-wavelength imaging of the Ca^2+^ indicator Fura-2 using a pulsed laser. 2P-imaging of Fura-2 in **Figures 1–2** of main text was performed in the single-wavelength (non-ratiometric) modality, to accommodate faster rates for imaging of the Ca^2+^ indicator and coincident photostimulation of the compounds at 2P. **Left:** Cultured HEK cells expressing mGlu_5_-eYFP receptors (yellow) and loaded with the ratiometric Ca^2+^ indicator Fura-2 AM (magenta). Numbers in parenthesis indicate wavelengths used for Fura-2 and eYFP excitation with pulsed and continuous-wave lasers, respectively. Arrowhead points at cell whose Ca^2+^ trace is shown in **B**. Scale bar is 50 μm. **Center**: Fura-2 was excited with the pulsed laser at different NIR wavelengths (760 to 820, in 20 nm steps) and a frame rate of 0.2 Hz. 2PE imaging wavelengths approximately correspond to 380–410 nm of 1PE imaging of Fura-2 with continuous-wave lasers. Therefore, increases in intracellular Ca^2+^ induced by bath-application of agonist (quisqualate, 3 μM) in cultured cells expressing mGlu_5_ are detected as decreases in Fura-2 fluorescence, consistent with its fluorescence emission at 380 nm (red arrow in right panel). **Right**: UV-vis spectra of Fura-2 excited at wavelengths indicated by horizontal axis, and in presence of increasing concentrations of free Ca^2+^ (modified from *leica-microsystems.com*). Red arrow illustrates direction of changes in Fura-2 fluorescence upon increases in free Ca^2+^ when the indicator is excited at 380 nm. **Panel B: Silencing of mGlu_5_ receptors using alloswitch in “all-2P experiments”**. Ca^2+^ trace in a cultured cell expressing mGlu_5_ and loaded with Fura-2 (excited with the pulsed laser at 780 nm, frame rate 0.2 Hz). Bath-application of agonist (quisqualate, 3 μM) at time indicated by blue arrowhead induces intracellular Ca^2+^ signaling (downward shift in 780 nm excited fluorescence of Fura-2), which is reverted to baseline after addition of alloswitch (1 μM, black arrowhead). **Panels C–E: “all-2P experiments” schematics (Figures 1–2) and increased baseline fluorescence of samples during photostimulation.** **Panel C**: Schematics of the imaging-photostimulation setup for “all-2P experiments” (shown in **Figures 1–2** of main text). Imaging of Fura-2 at 2PE is interleaved with photostimulations of alloswitches at 2PE, and performed at the same wavelength but different power values of the pulsed laser, by raster-scanning with an inverted confocal microscope. For 2PE imaging of Fura-2, low-power (3 mW) and slow frame rates (0.2 Hz) are used, whereas photostimulation of alloswitches is achieved by consecutive raster scans at the same wavelength (Δλ = 760, 780, 800, or 820 nm) but high power (12 mW) and maximum frame rate. The top trace is an example of a Ca^2+^ imaging session in cultured cells, and zoom-in of the first part of the trace shows sequential imaging scans for visualization of Ca^2+^ dynamics through 2P-imaging of Fura-2. Zoom-in of the photostimulation period (red box) indicates that photostimulation was accomplished by 13 sequential scans at maximum speed and was followed by one imaging scan at the same wavelength but low power; this cycle of “photostimulation+imaging” was repeated 36 times for calculation of oscillatory frequencies during photostimulation. Laser power during imaging scans was set to minimal values allowing satisfactory visualization of Fura-2 (~3 mW on sample for all wavelengths used), with the rationale of minimizing unwanted 2PE of alloswitch molecules due to imaging of Fura-2. **Panels D–E**: In “all-2P experiments”, steady increases in the basal fluorescence of all samples were observed during photostimulation periods (light-red boxes). These increases reflected increased basal fluorescence of the sample rather than real decreases in intracellular Ca^2+^, and were due to an arbitrarily increased light power of imaging scans during photostimulation done by the microscope software (see **Methods 5.2.2** for details). To verify this, low-power (3 mW) scans for 2P-imaging of Fura-2 in cultured HEK cells (**D**) or for visualization of a fluorescent plastic slide by Chroma (E) were combined with mock high-power (20 mW) 2PE scans during photostimulation periods. Importantly, the increased fluorescence during photostimulation was compatible with the detection of Ca^2+^ oscillations, as illustrated in **panel D**. These increases in baseline fluorescence were not *post-hoc* corrected in any of the cell traces shown throughout the paper. Agonist for mGlu_5_ activation is quisqualate (3 μM, blue triangle); fluorescence intensity is shown in arbitrary units over frame number.

**Figure S2 (related to Figure 2):**
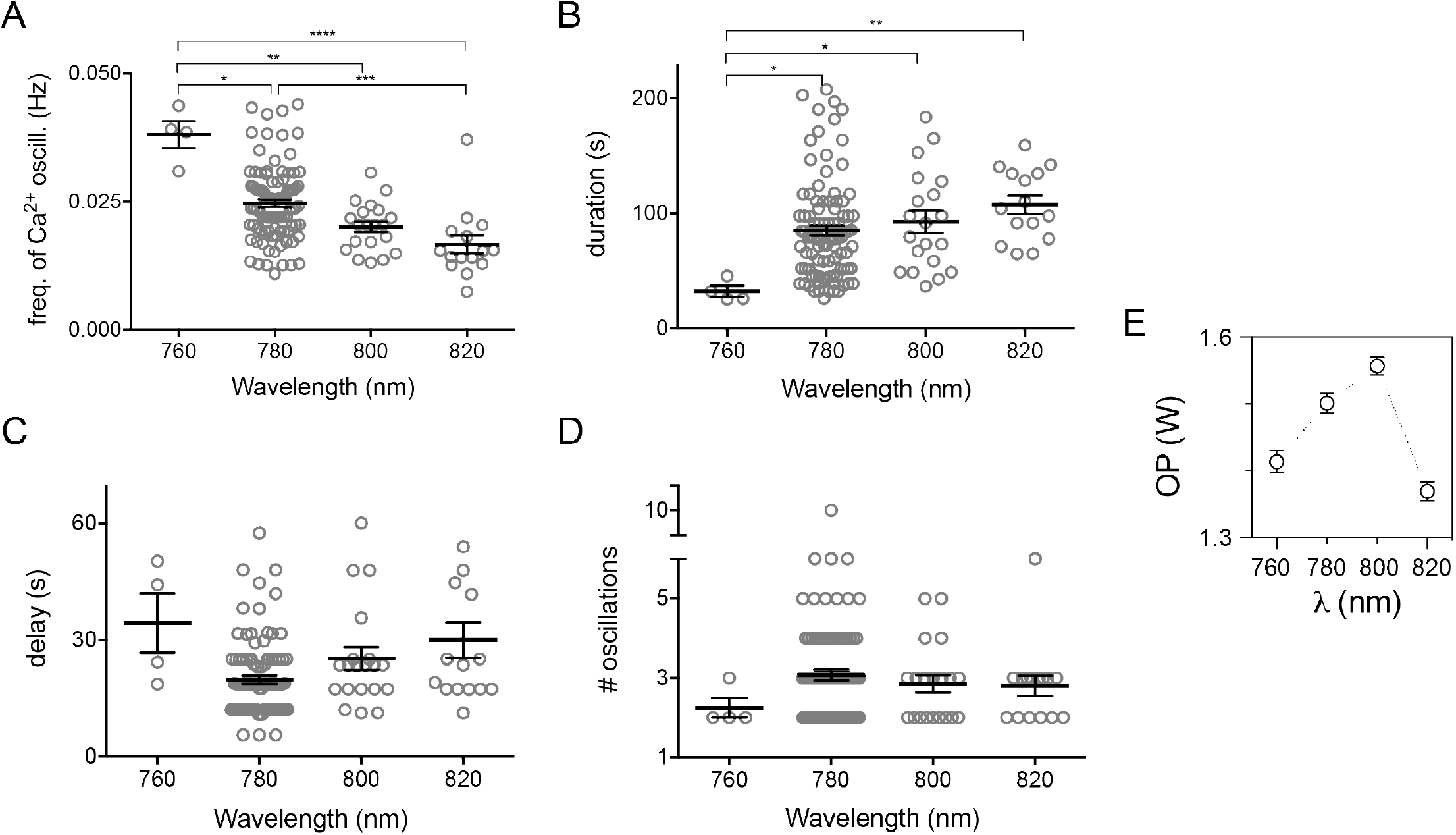
Optimization of 2PE conditions for Fura-2 fluorescence imaging and alloswitch isomerization. **Panels A–D**: Full dataset and statistics for functional efficiency of 2PE of alloswitch at different infrared wavelengths (Figure 2 of main text). In brief, Fura-2 fluorescence was used to monitor Ca^2+^ oscillations in cultured HEK cells expressing mGlu_5_-eYFP receptors. Oscillations were photoinduced by 2PE (13 scans, 12 mW laser power) of alloswitch (1 μM) in bath along with agonist (quisqualate, 3 μM) at wavelengths indicated by horizontal axis. Graphs illustrate functional efficiency of alloswitch isomerization at 2PE, expressed as the frequency of photoinduced Ca^2+^ oscillations **(A)**, duration of the oscillatory behavior **(B)**, time for onset of first Ca^2+^ peak after photostimulation start (delay, **C**), and average number of oscillations counted during 2PE (# oscillations, **D**). Data represented as mean ± s.e.m., n = 4–95 cells. * *p* < 0.05, ** *p* < 0.01, *** *p* < 0.001, **** *p* < 0.0001, using Dunn’s multiple comparison test after the Kruskal–Wallis test. **Panel E**: Output power of the pulsed laser at the wavelengths used for photostimulation of alloswitch in **A–D**.

**Figure S3 (related to Figure 1):**
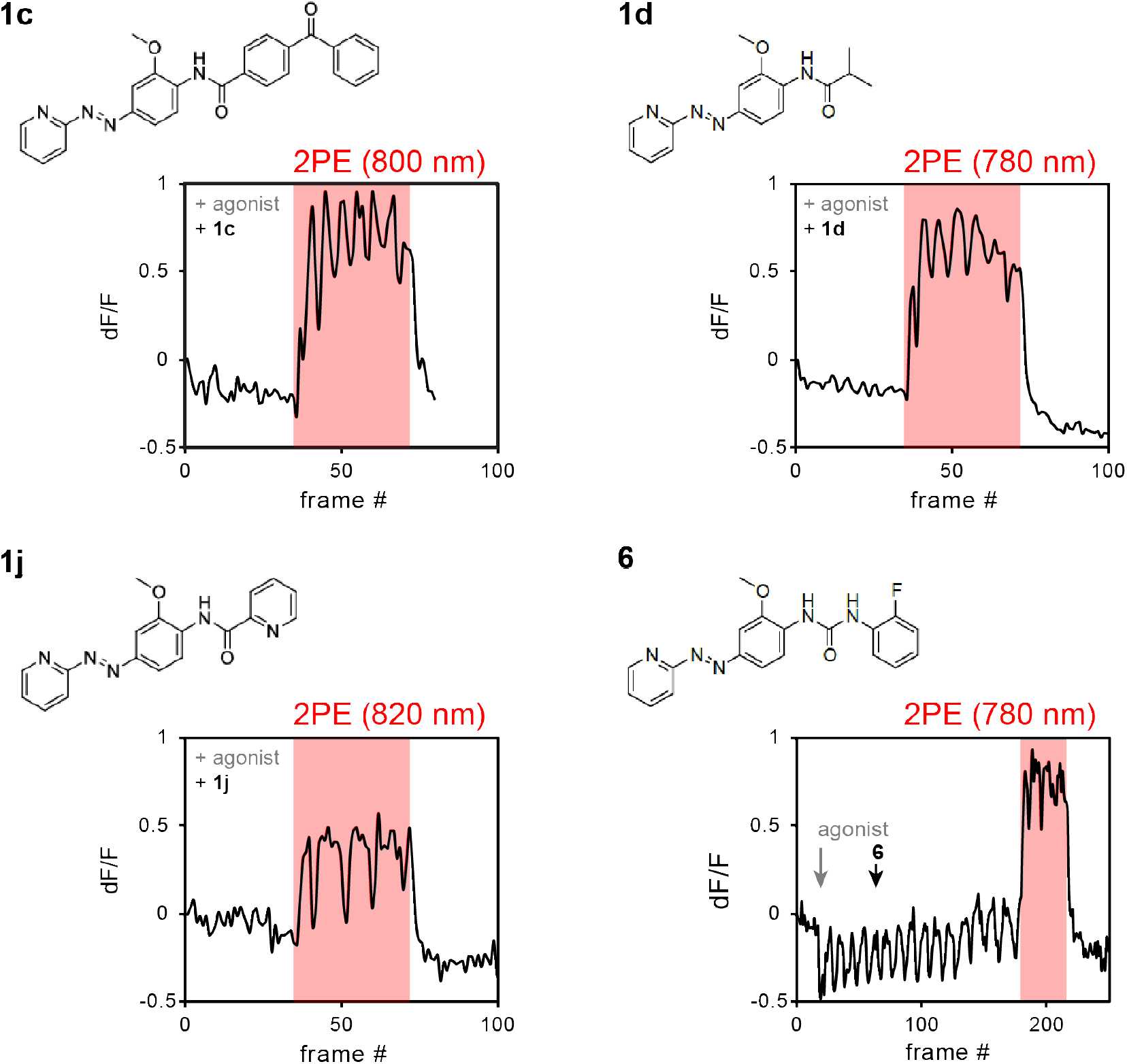
“All-2P experiments”, structures of alloswitches used, and example traces of their 2PE. Schematics of imaging-photostimulation setup for “all-2P experiments” can be found in **Figure S1C.** Compounds **1c-d, 1j, 6** are alloswitch analogs used for “all-2P experiments” other than alloswitch (names correspond to compounds published in Gómez-Santacana *et al.,* 2017; parent alloswitch structure is shown in **Figure 1** of main text). Compound structures are shown, along with example traces of 2PE at wavelengths indicated above trace boxes. Traces show changes in Fura-2 fluorescence (dF/F) in mGlu_5_-eYFP expressing HEK cells, supplemented with agonist (quisqualate, 3 μM) and alloswitch analogs (1 μM) prior to experiment (except for compound **6** where additions are indicated by grey and black arrows, respectively). Compounds were selected based on structural variations aimed at improving the performance of alloswitch at 2PE. In particular, we focused on replacing the 2-chlorophenyl group of alloswitch with an aliphatic amide (1d) or stronger electron acceptors (benzophenone in **1c**, pyridine in **1j**, urea and 2-flurobenzene in **6**), thus increasing internal charge transfer upon electron excitation (push-pull character) of the phenylazopyridine, and, as such, two-photon absorption^1^. Consistent with an increased push-pull character, compounds **1d** and **6** display, respectively, improved potency and red-shifted spectrum compared to alloswitch^2^. Effective photoisomerization during 2PE at indicated wavelengths was observed for all alloswitch analogs tested except compound **6**, that failed to silence Ca^2+^ oscillations induced by agonist at the concentration used (1 μM) and no inferences about his 2PE character were possible. Extensive wavelength-dependence characterization (as performed for the parent molecule in **Figure 2** of main text) is required to determine relative performance of alloswitch analogs at 2PE.

**Figure S4 (related to Figures 3 and 4):**
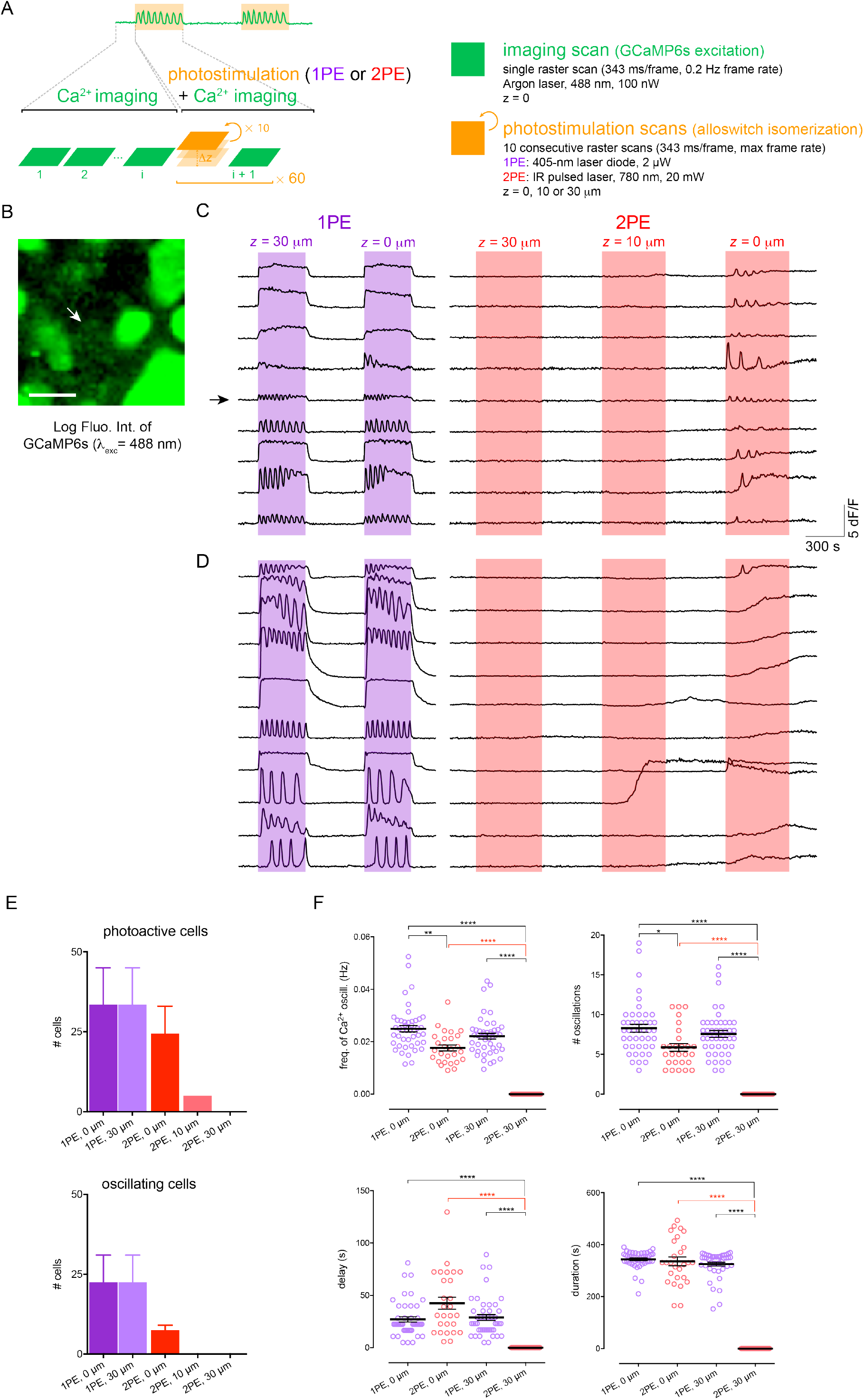
Panels A–D: Schematics of photostimulation experiments at different axial distances in cultured cells (Figures 3 and 4). **Panel A**: Schematics of setup used to combine imaging of intracellular Ca^2+^ (green) with 1PE or 2PE photostimulation (orange) of alloswitches, by raster-scanning with an inverted confocal microscope. The top trace corresponds to a Ca^2+^ imaging session in cultured cells, and zoom-in of the first part of the trace shows sequential imaging scans for visualization of Ca^2+^ through GCaMP6s fluorescence. Zoom-in of the photostimulation period (light orange box) indicates that photostimulation was accomplished by 10 scans at maximum speed followed by one imaging scan, and this cycle photostimulation-imaging was repeated 60 times. The 10 photostimulation scans were done at one axial position (Δz = 30, 10 or 0 μm) above the imaging plane (z = 0). Details of scanning and laser settings, as well as axial positions are listed on the right. **Panel B**: Cultured HEK cells co-expressing mGlu_5_ receptors and GCaMP6s (excited at 488 nm, 100 nW; shown as logarithm of fluorescence intensity for better visualization) for Ca^2+^ imaging at 1PE. Arrow indicates cell shown by arrow in C, scale bar is 20 μm. **Panels C-D**: Example traces of cultured HEK cells where changes in intracellular Ca^2+^ were monitored by imaging GCaMP6s in the presence of mGlu_5_ agonist (quisqualate, 3 μM) and alloswitch (1 μM), and in the absence or presence of photostimulation of alloswitch at 1PE (violet boxes; 405 nm, 2 μW, repetition rate 0.17 Hz) or 2PE (red boxes; 780 nm, 20 mW, repetition rate 0.12 Hz) performed at axial distances indicated above colored boxes. Cells oscillating when photostimulated at 2PE at the imaging plane (z = 0) are shown in **C**; examples of cells showing Ca^2+^ events other than oscillations under 2PE (used to compute % of photostimulated cells in **Figure 4, panel D**) are shown in **D**. Ca^2+^ traces are shown as fluorescence changes relative to first frame (dF/F) of GCaMP6s (excited at 488 nm, 100 nW). **Panel E: Average number of cells photoactive or oscillating at 1PE and 2PE, and different axial positions of photostimulation (related to cell percentages presented in Figure 4, panel D).** In brief, changes in intracellular Ca^2+^ were monitored by imaging GCaMP6s in the presence of mGlu_5_ agonist (quisqualate, 3 μM) and alloswitch (1 μM), and in the absence or presence of photostimulation of alloswitch at 1PE (violet boxes; 405 nm, 2 μW, repetition rate 0.17 Hz) or 2PE (red boxes; 780 nm, 20 mW, repetition rate 0.12 Hz) performed at axial distances indicated by horizontal axes. Photoactive cells (**top**) comprise cells responding to photostimulation with Ca^2+^ oscillations (oscillating cells, bottom) or with other types of intracellular Ca^2+^ events (examples of cells photoactive under 2PE but not oscillating can be viewed in panel D). Ca^2+^ events observed were either steady Ca^2+^ increases or peak-and-plateau responses. Cell classification was conducted as detailed in ***Methods, Section 6.2**.* The relative abundance of Ca^2+^ responses other than oscillations in cultured cells is not surprising, given the different amounts of receptor and G-protein subunit expression have been previously demonstrate to alter the phenotype of intracellular Ca^2+^ events in response to activation of mGlu_5_ receptors^3,4^. The total number of cells responsive to 1PE at the imaging plane (‘1 PE, 0 μm’, **top**) was used to calculate the percentage of photoactive and oscillating cells shown in **Figure 4, panel D** of main text. Data are average number of cells for each condition; n = 67 cells from 2 independent experiments. **Panel F: Full datasets and statistics for photoactivation experiments at 1PE or 2PE and different axial distances in cultured cells (shown in **Figure 4, panel E** of main text)**. Briefly, cultured HEK cells co-expressing mGlu_5_ receptors and GCaMP6s for Ca^2+^ recordings were imaged as detailed in **Figure 4**. The oscillatory behavior in response to agonist (quisqualate, 3 μM) was monitored during silencing of receptors with alloswitch (1 μM) and coincident photoactivation at 1PE (violet) or 2PE (red) done at different axial distances (indicated in horizontal axes) from the cell imaging plane (0 μm). Photoactivation is quantified as frequency and total number of calcium oscillations, time for the peak of the first oscillation observed after light onset (delay), and overall duration of the oscillatory behavior (duration). Lines and error bars represent mean ± s.e.m. plotted above individual cells (empty circles). * *p* < 0.05, ** *p* < 0.01, *** *p* < 0.001, **** *p* < 0.0001, using Dunn’s correction for multiple comparisons after the Kruskal-Wallis test. D’Agostino and Pearson omnibus normality test was significant only for the dataset ‘2PE, 0 μm’ in all variables measured except for the delay. For 1PE conditions, n = 45 from 2 independent experiments; for 2PE, n = 27 from 3 independent experiments.

**Figure S5 (related to Figure 5):**
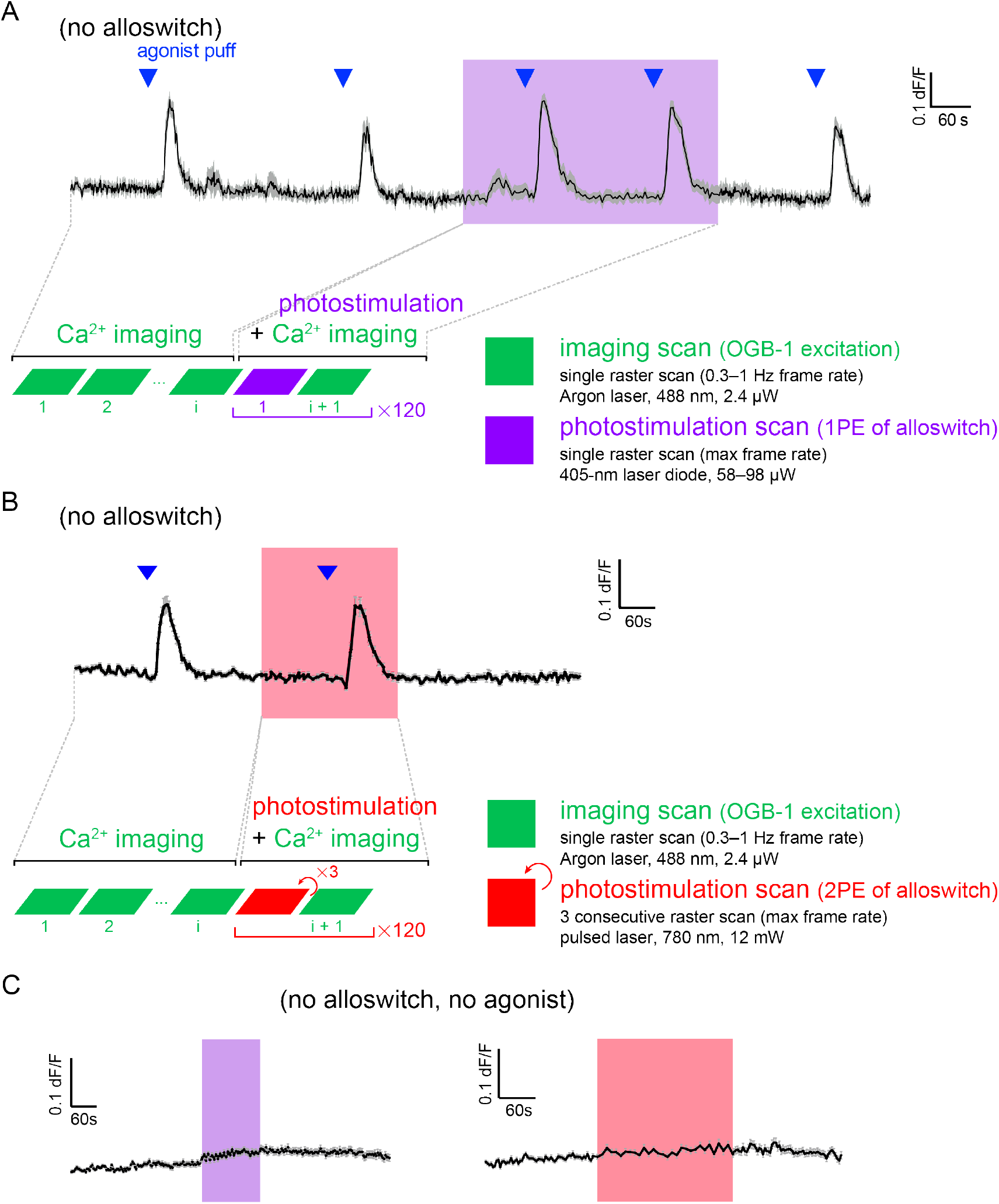
Panels A–B: Schematics of imaging-photostimulation cycles in rat acute brain slices and control experiments. **Panel A**: Schematics at the **bottom of the Figure** illustrate the setup used to monitor intracellular Ca^2+^ in acute brain slices experiments where OGB-1 imaging (green) was combined with photostimulation at 1PE (violet) by raster-scanning with an inverted confocal microscope. The photostimulation period (light violet box in top trace) is composed of 120 repetitions of: one raster scan of the 405 nm laser diode, followed by one 488 nm imaging scan for visualization of OGB-1. Details of scanning and laser settings used across experiments are listed on the right. **Top of Figure** shows the average Ca^2+^ trace (mean ± s.e.m., n = 16 cells; 1 P9 rat brain slice) of a control experiment for photostimulation at 1PE in hippocampal slices, performed in the absence of alloswitch. 1PE (violet box) is done at a laser power (98 μW, pixel dwell time = 2.5 μs) almost twice the power used for successful photostimulation of alloswitch in **Figure 5, panel D** (58 μW). Intracellular Ca^2+^ is shown as changes in OGB-1 fluorescence (dF/F) over time. Activation of endogenous mGlu_5_ receptors is done by pressure-ejection of agonist as in experiments in **Figure 5** (DHPG, 1 mM; 800 hPa, 1 s), before, during and after photostimulation. **Panel B**: Average trace (n=112 cells, 1 animal) of DHPG (1mM; 500 hPa, 1 s)-mediated Ca^2+^ response at both dark and 2PE (25 mW, 0.3 Hz) in the absence of alloswitch and corresponding schematics of how imaging and photostimulation have been performed. **Panel C: Control experiments to determine the effect of photostimulation alone on intracellular Ca^2+^ in slices.** **Left**: Average trace (n=238 cells, 3 animals) of 1P light control experiments. Acute brain slices were perfused with normal ACSF and illuminated with the 405 nm laser diode (violet box) at 58 μW light power and 0.4 Hz illumination rate. **Right**: Average trace (n = 171 cells, 2 animals) of 2PE control experiments on acute brain slices. 780 nm (red box) at 25 mW, 0.3 Hz.

**Figure S6 (related to Figures 5-7):**
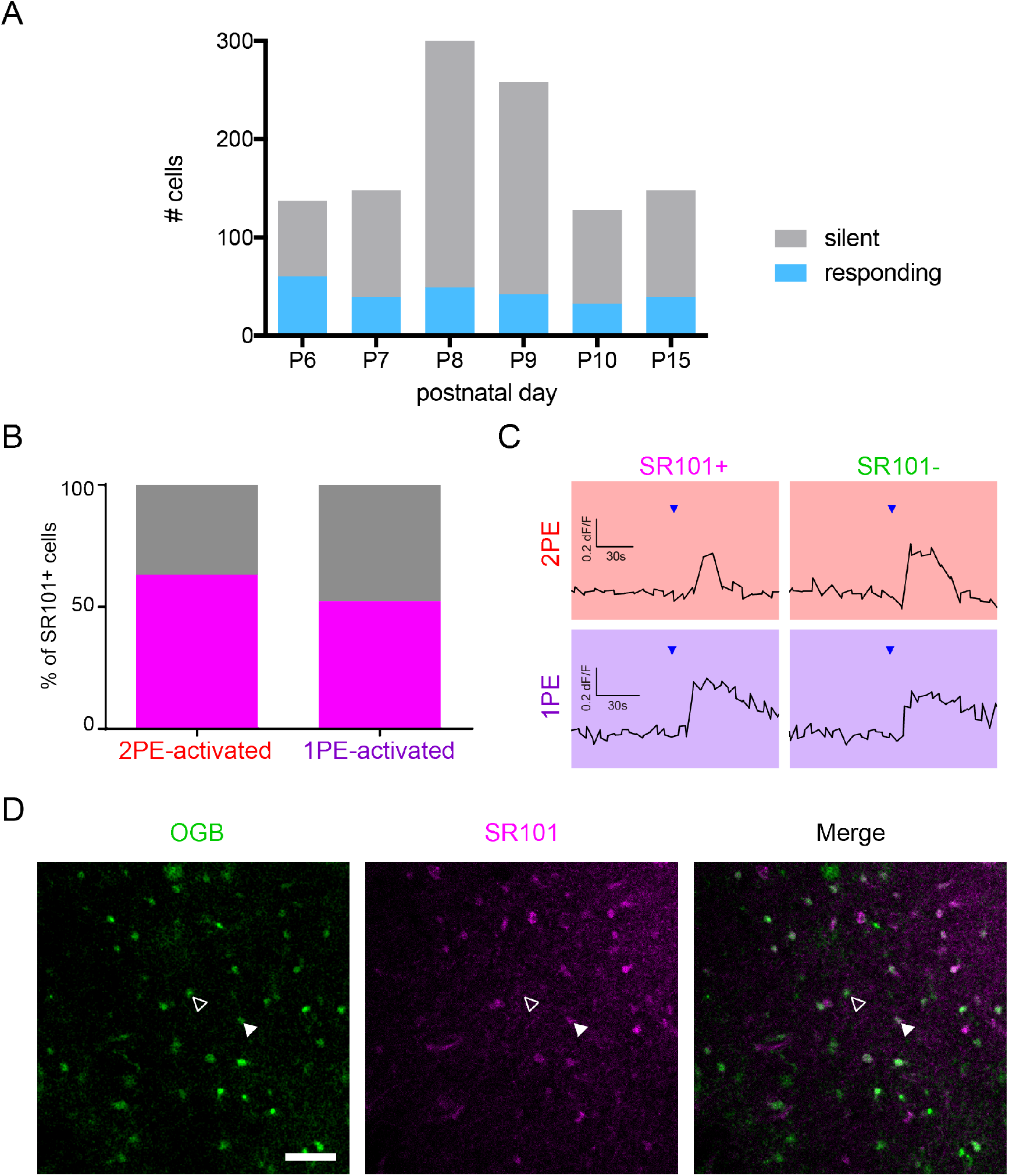
Panel A: Outline of experiments performed at different postnatal ages in acute rodent brain slices. Percentage of cells in *naïve* slices obtained from animals of different ages (P6–P15) responding to local puffing of mGlu_5_ agonist (DHPG, 1 mM; 500–800 hPa, 1 s). Exploratory experiments were performed at indicated ages to identify optimal experimental conditions; P8-P9 rats were selected for further experiments because of optimal OGB-1 labeling (in our hands, labeling was poorer after P10). The increased number in labeled cells could also be due to better survival of cells after sectioning at these ages, which might account for the relatively constant number of responding cells across ages. **Panels B–D: Proportion of SR101+ cells photoactivated in hippocampal CA1 *stratum radiatum* as in Figure 6 and 7 of main text.** **Panel B:** Percentage of SR101+ cells among all cells analyzed in **Figure 7** of the main text (n=114 cells from 3 animals). For 2PE, 63 ± 15% of the responding cells were SR101+ and for 1PE, 53 ± 14% (Unpaired t-test, p=0.633). **Panel C**: Example traces of SR101+ cell and SR101^-^ cell during 2PE (red boxes) and 1PE (violet boxes) are presented as changes in OGB fluorescence (dF/F) over time. Agonist (blue arrowheads, 1 mM DHPG) was locally ejected at 500 hPa during 1 second. The average amplitude of calcium responses (n = 114 cells, 3 animals) was slightly higher (p = 0.035, unpaired t-test) in astrocytes (SR101+) compared to neurons (SR101^-^) during photoactivation of alloswitch at 1PE but not 2PE (p = 0.091, unpaired t-test), probably reflecting a more extended volumetric spread of astrocytes compared to neurons. **Panel D**: Cells in *stratum radiatum* labeled with OGB-1 (left), astrocytic marker SR101(middle), and the two channels merged (right). One photoactivated SR101+ (white arrowhead) and one SR101^-^ (empty arrowhead) cell are indicated. Scale bar is 50 μm.

**Figure S7 (related to Figure 7):**
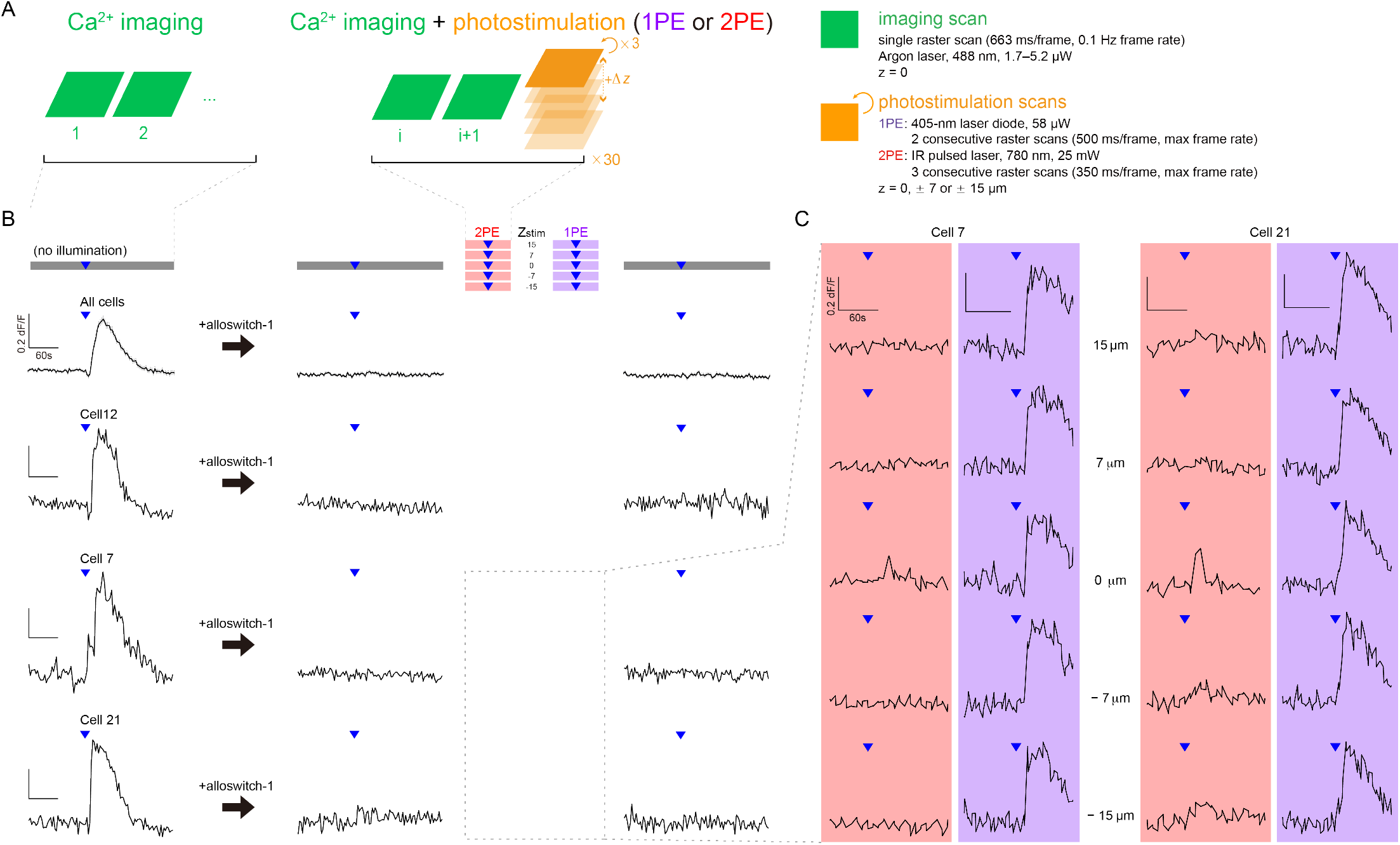
Inhibition of mGlu_5_ activity by alloswitch was assessed throughout experiments shown in Figure 7 of the main text. **Panel A**: Schematics for experiments in slices as in **Figure 7**, illustrating how Ca^2+^ imaging and photostimulation of alloswitch were performed along with changes in axial position for photostimulation (Δ*z*). **Panel B**: DHPG-mediated (blue triangles) Ca^2+^ responses were monitored in the absence of illumination (grey bars) before and after applying alloswitch, and during photostimulation (colored bars). After consecutive stimulations with 2PE (red) and 1PE (violet) at 5 different axial distances (z_stim_) from the cell imaging plane, the inhibition exerted by *trans*-alloswitch was maintained in the absence of illumination. Top trace corresponds to averaged recordings from all cells analyzed in **Figure 7 C-D** of the main text; n=114 cells from 3 animals. Cell 12 is the same cell shown in **Figure 7 B**. Cell 7 and Cell 21 are additional examples of cells responding with Ca^2+^ peaks to agonist ejection (blue triangles) before addition of alloswitch (left traces), and demonstrating complete silencing by *trans*-alloswitch before (middle traces) and after (right traces) light stimulation protocols at 5 different axial distances (z_stim_) and two light conditions (red/violet). **Panel C**: Ca^2+^ responses of Cell 7 and Cell 21 from **A** during 2PE (red) and 1PE (violet) at different axial distances (from the top, z_stim_= +15, +7, 0, −7, −15 μm) show a reversible control of mGlu_5_-mediated response by photoswitching, which is axially restricted for 2PE but not for 1PE.

### Supplemental Movies

**Movie S1 (related to Fig.3):**

**Photorescue of mGlu_5_ receptors by 1PE and 2PE in HEK cells.**

Video shows GCaMP6s fluorescence in cultured HEK cells co-transfected with mGlu_5_ receptors as in **Figure 3**. Cell in the middle corresponds to cell indicated by arrow in **Figure 3B** and Ca^2+^ trace in **Figure 3C**, and responds with Ca^2+^ oscillations in all conditions. Intracellular events are recorded at conditions indicated by labels (“agonist” = 3 μM quisqualate; ‘+ alloswitch/no light’ = 1 μM alloswitch supplemented in bath in addition to agonist; ‘1PE’ = 405 nm laser diode, 2 μW; ‘2PE’ = NIR pulsed laser, 780 nm, 20 mW; see also **Methods, Section 5.2**). Photostimulation periods start on frame 6 after label, and last 60 frames each. GCaMP6s fluorescence is excited by raster scanning with a 488 nm Argon laser (100 nW, 343 ms/frame). Frame rate is 0.2 Hz during imaging alone, 0.17 Hz during 1P photostimulation, and 0.12 Hz during 2P photostimulation. Picture size is 62×62 μm.

**Movie S2 (related to Fig.4):**

**Axial-plane specific photorescue of mGlu_5_ receptors by 2PE but not 1PE in HEK cells.**

Video of GCaMP6s fluorescence in cultured HEK cells co-transfected with mGlu_5_ receptors. Photocontrol of cell Ca^2+^ occurs regardless of the axial distance for 1PE of alloswitch, and is axial-plane selective for 2PE. The field of view corresponds to image and cell Ca^2+^ trace in **Figure 4B-C** in main text. Ca^2+^ events are recorded in different conditions indicated by labels (‘agonist’ = 3 μM quisqualate; ‘+ alloswitch/no light’ = 1 μM alloswitch supplemented in bath in addition to agonist; ‘1PE’ = 405 nm laser diode, 2 μW; ‘2PE’ = NIR pulsed laser, 780 nm, 20 mW; ‘light OFF’ = end of photostimulation; see also **Methods, Section 5.2**). Axial position of photostimulation (z in main figures) is indicated between asterisks, and is 30 or 10 μm above cells, or corresponds to the cell imaging plane (0 μm). GCaMP6s fluorescence is excited by raster scanning with a 488 nm Argon laser (100 nW, 343 ms/frame). Frame rate is 0.2 Hz during imaging alone, 0.17 Hz during 1P photostimulation, and 0.12 Hz during 2P photostimulation. Picture size is 129×129 μm.

**Movie S3 (related to Fig.5):**

**Silencing and photorescue of mGlu_5_ receptors by alloswitch and 1PE in acute brain slices.**

Video of OGB-1 fluorescence in a P9 rat acute brain slice, corresponding to images and Ca^2+^ traces in **Figure 5B-D** in main text. Intracellular Ca^2+^ events are recorded at baseline and different drug and photostimulation conditions indicated by labels (‘baseline’ = aCSF containing TTX/LY367385; ‘+ alloswitch/no light’ = 10 μM alloswitch supplemented in recirculating aCSF; white dot indicates site and time of agonist pressure-ejection: DHPG, 1 mM and 800 hPa, 1 s; ‘1PE’ = 405 nm laser diode, 58 μW, 120 raster scans at 0.3 Hz; see also **Methods, Section 5.2**). OGB-1 fluorescence is excited by raster scanning with a 488 nm Argon laser (2.4 μW, 1 Hz frame rate). Frame rate for OGB-1 is 0.3 Hz during photostimulation. Picture size is 350×350 μm.

**Movie S4 (related to Fig.6):**

**Photorescue of mGlu_5_ receptors by 2PE in acute brain slices.**

Video of OGB-1 fluorescence in a P10 rat acute brain slice, corresponding to Ca^2+^ traces in **Figure 6B**. Intracellular Ca^2+^ events are recorded in the dark and under photostimulation conditions indicated by labels (‘+ alloswitch/no light’ = 10 μM alloswitch supplemented in bubbling reservoir; ‘2PE’ = 780 nm pulsed laser, 25 mW, 120 raster scans at 0.3 Hz; see also **Methods, Section 5.2**). The white dot indicates the site and time of agonist pressure-ejection (DHPG, 1 mM; 500 hPa, 1 s). OGB-1 fluorescence is excited by raster scanning with a 488 nm Argon laser (2.4 μW, 1 Hz frame rate). Frame rate for OGB-1 is 0.2 Hz during photostimulation. Picture size is 310×310 μm.

## References

1 Kim, H. & Kim, J. S. A guide to genome engineering with programmable nucleases. Nat Rev Genet 15, 321–334, doi:10.1038/nrg3686 (2014).

2 Ghildiyal, M. & Zamore, P. D. Small silencing RNAs: an expanding universe. Nat Rev Genet 10, 94–108, doi:10.1038/nrg2504 (2009).

3 Behar, M., Barken, D., Werner, S. L. & Hoffmann, A. The dynamics of signaling as a pharmacological target. Cell 155, 448–461, doi:10.1016/j.cell.2013.09.018 (2013).

4 Reiner, A. & Isacoff, E. Y. Tethered ligands reveal glutamate receptor desensitization depends on subunit occupancy. Nat Chem Biol 10, 273–280, doi:10.1038/nchembio.1458 (2014).

5 Velema, W. A., Szymanski, W. & Feringa, B. L. Photopharmacology: beyond proof of principle. J Am Chem Soc 136, 2178–2191, doi:10.1021/ja413063e (2014).

6 Schaefer, K. A. et al. Unexpected mutations after CRISPR-Cas9 editing in vivo. Nat Methods 14, 547–548, doi:10.1038/nmeth.4293 (2017).

7 Kienzler, M. A. & Isacoff, E. Y. Precise modulation of neuronal activity with synthetic photoswitchable ligands. Curr Opin Neurobiol 45, 202–209, doi:10.1016/j.conb.2017.05.021 (2017).

8 Miesenbock, G. Optogenetic control of cells and circuits. Annu Rev Cell Dev Biol 27, 731–758, doi:10.1146/annurev-cellbio-100109-104051 (2011).

9 Rost, B. R., Schneider-Warme, F., Schmitz, D. & Hegemann, P. Optogenetic Tools for Subcellular Applications in Neuroscience. Neuron 96, 572–603, doi:10.1016/j.neuron.2017.09.047 (2017).

10 Miyashita, T., Shao, Y. R., Chung, J., Pourzia, O. & Feldman, D. E. Long-term channelrhodopsin-2 (ChR2) expression can induce abnormal axonal morphology and targeting in cerebral cortex. Front Neural Circuits 7, 8, doi:10.3389/fncir.2013.00008 (2013).

11 Wu, W. et al. Increased threshold of short-latency motor evoked potentials in transgenic mice expressing Channelrhodopsin-2. PLoS One 12, e0178803, doi:10.1371/journal.pone.0178803 (2017).

12 Volgraf, M. et al. Reversibly caged glutamate: a photochromic agonist of ionotropic glutamate receptors. J Am Chem Soc 129, 260–261, doi:10.1021/ja067269o (2007).

13 Stawski, P., Sumser, M. & Trauner, D. A photochromic agonist of AMPA receptors. Angew Chem Int Ed Engl 51, 5748–5751, doi:10.1002/anie.201109265 (2012).

14 Barber, D. M. et al. Optical control of AMPA receptors using a photoswitchable quinoxaline-2,3-dione antagonist. Chem Sci 8, 611–615, doi:10.1039/c6sc01621a (2017).

15 Laprell, L. et al. Optical control of NMDA receptors with a diffusible photoswitch. Nat Commun 6, 8076, doi:10.1038/ncomms9076 (2015).

16 Pittolo, S. et al. An allosteric modulator to control endogenous G protein-coupled receptors with light. Nat Chem Biol 10, 813–815, doi:10.1038/nchembio.1612 (2014).

17 Rovira, X. et al. OptoGluNAM4.1, a Photoswitchable Allosteric Antagonist for Real-Time Control of mGlu4 Receptor Activity. Cell Chem Biol 23, 929–934, doi:10.1016/j.chembiol.2016.06.013 (2016).

18 Banghart, M. R. & Sabatini, B. L. Photoactivatable neuropeptides for spatiotemporally precise delivery of opioids in neural tissue. Neuron 73, 249–259, doi:10.1016/j.neuron.2011.11.016 (2012).

19 Schonberger, M. & Trauner, D. A photochromic agonist for mu-opioid receptors. Angew Chem Int Ed Engl 53, 3264–3267, doi:10.1002/anie.201309633 (2014).

20 Stein, M. et al. Azo-propofols: photochromic potentiators of GABA(A) receptors. Angew Chem Int Ed Engl 51, 10500–10504, doi:10.1002/anie.201205475 (2012).

21 Damijonaitis, A. et al. AzoCholine Enables Optical Control of Alpha 7 Nicotinic Acetylcholine Receptors in Neural Networks. ACS Chem Neurosci 6, 701–707, doi:10.1021/acschemneuro.5b00030 (2015).

22 Bahamonde, M. I. et al. Photomodulation of G protein-coupled adenosine receptors by a novel light-switchable ligand. Bioconjug Chem 25, 1847–1854, doi:10.1021/bc5003373 (2014).

23 Denk, W., Strickler, J. H. & Webb, W. W. Two-photon laser scanning fluorescence microscopy. Science 248, 73–76 (1990).

24 Svoboda, K. & Yasuda, R. Principles of two-photon excitation microscopy and its applications to neuroscience. Neuron 50, 823–839, doi:10.1016/j.neuron.2006.05.019 (2006).

25 Packer, A. M., Roska, B. & Hausser, M. Targeting neurons and photons for optogenetics. Nat Neurosci 16, 805–815, doi:10.1038/nn.3427 (2013).

26 Izquierdo-Serra, M. et al. Two-photon neuronal and astrocytic stimulation with azobenzene-based photoswitches. J Am Chem Soc 136, 8693–8701, doi:10.1021/ja5026326 (2014).

27 Carroll, E. C. et al. Two-photon brightness of azobenzene photoswitches designed for glutamate receptor optogenetics. Proc Natl Acad Sci U S A 112, E776–785, doi:10.1073/pnas.1416942112 (2015).

28 Gomez-Santacana, X. et al. Illuminating Phenylazopyridines To Photoswitch Metabotropic Glutamate Receptors: From the Flask to the Animals. ACS Cent Sci 3, 81–91, doi:10.1021/acscentsci.6b00353 (2017).

29 Huber, K. M., Kayser, M. S. & Bear, M. F. Role for rapid dendritic protein synthesis in hippocampal mGluR-dependent long-term depression. Science 288, 1254–1257 (2000).

30 Wang, X. et al. Altered mGluR5-Homer scaffolds and corticostriatal connectivity in a Shank3 complete knockout model of autism. Nat Commun 7, 11459, doi:10.1038/ncomms11459 (2016).

31 Aloisi, E. et al. Altered surface mGluR5 dynamics provoke synaptic NMDAR dysfunction and cognitive defects in Fmr1 knockout mice. Nat Commun 8, 1103, doi:10.1038/s41467-017-01191-2 (2017).

32 Matosin, N., Fernandez-Enright, F., Lum, J. S. & Newell, K. A. Shifting towards a model of mGluR5 dysregulation in schizophrenia: Consequences for future schizophrenia treatment. Neuropharmacology 115, 73–91, doi:10.1016/j.neuropharm.2015.08.003 (2017).

33 Shin, S. et al. mGluR5 in the nucleus accumbens is critical for promoting resilience to chronic stress. Nat Neurosci 18, 1017–1024, doi:10.1038/nn.4028 (2015).

34 Haas, L. T. et al. Silent Allosteric Modulation of mGluR5 Maintains Glutamate Signaling while Rescuing Alzheimer’s Mouse Phenotypes. Cell Rep 20, 76–88, doi:10.1016/j.celrep.2017.06.023 (2017).

35 Ferreira, D. G. et al. alpha-synuclein interacts with PrPC to induce cognitive impairment through mGluR5 and NMDAR2B. Nat Neurosci 20, 1569–1579, doi:10.1038/nn.4648 (2017).

36 Xu, C. & Zipfel, W. R. Multiphoton excitation of fluorescent probes. Cold Spring Harb Protoc 2015, 250–258, doi:10.1101/pdb.top086116 (2015).

37 Chen, T. W. et al. Ultrasensitive fluorescent proteins for imaging neuronal activity. Nature 499, 295–300, doi:10.1038/nature12354 (2013).

38 Nash, M. S. et al. Determinants of metabotropic glutamate receptor-5-mediated Ca2+ and inositol 1,4,5-trisphosphate oscillation frequency. Receptor density versus agonist concentration. J Biol Chem 277, 35947–35960, doi:10.1074/jbc.M205622200 (2002).

39 Sun, W. et al. Glutamate-dependent neuroglial calcium signaling differs between young and adult brain. Science 339, 197–200, doi:10.1126/science.1226740 (2013).

40 Romano, C. et al. Distribution of metabotropic glutamate receptor mGluR5 immunoreactivity in rat brain. J Comp Neurol 355, 455–469, doi:10.1002/cne.903550310 (1995).

41 Rossi, A. et al. Genetic compensation induced by deleterious mutations but not gene knockdowns. Nature 524, 230–233, doi:10.1038/nature14580 (2015).

42 El-Brolosy, M. A. & Stainier, D. Y. R. Genetic compensation: A phenomenon in search of mechanisms. PLoS Genet 13, e1006780, doi:10.1371/journal.pgen.1006780 (2017).

43 Kakkar, A., Traverso, G., Farokhzad, O. C., Weissleder, R. & Langer, R. Evolution of macromolecular complexity in drug delivery systems. Nature Reviews Chemistry 1, 0063, doi:10.1038/s41570-017-0063 (2017).

44 Mittmann, W. et al. Two-photon calcium imaging of evoked activity from L5 somatosensory neurons in vivo. Nat Neurosci 14, 1089–1093, doi:10.1038/nn.2879 (2011).

45 Nikolenko, V. et al. SLM Microscopy: Scanless Two-Photon Imaging and Photostimulation with Spatial Light Modulators. Front Neural Circuits 2, 5, doi:10.3389/neuro.04.005.2008 (2008).

46 Anselmi, F., Ventalon, C., Begue, A., Ogden, D. & Emiliani, V. Three-dimensional imaging and photostimulation by remote-focusing and holographic light patterning. Proc Natl Acad Sci U S A 108, 19504–19509, doi:10.1073/pnas.1109111108 (2011).

47 Papagiakoumou, E. et al. Scanless two-photon excitation of channelrhodopsin-2. Nat Methods 7, 848–854, doi:10.1038/nmeth.1505 (2010).

48 Dal Maschio, M., Donovan, J. C., Helmbrecht, T. O. & Baier, H. Linking Neurons to Network Function and Behavior by Two-Photon Holographic Optogenetics and Volumetric Imaging. Neuron 94, 774–789 e775, doi:10.1016/j.neuron.2017.04.034 (2017).

49 Prevedel, R. et al. Fast volumetric calcium imaging across multiple cortical layers using sculpted light. Nat Methods 13, 1021–1028, doi:10.1038/nmeth.4040 (2016).

50 Forli, A. et al. Two-Photon Bidirectional Control and Imaging of Neuronal Excitability with High Spatial Resolution In Vivo. Cell Rep 22, 3087–3098, doi:10.1016/j.celrep.2018.02.063 (2018).

51 Shemesh, O. A. et al. Temporally precise single-cell-resolution optogenetics. Nat Neurosci 20, 1796–1806, doi:10.1038/s41593-017-0018-8 (2017).

52 Mardinly, A. R. et al. Precise multimodal optical control of neural ensemble activity. Nat Neurosci 21, 881–893, doi:10.1038/s41593-018-0139-8 (2018).

53 Ellis-Davies, G. C. Two-photon microscopy for chemical neuroscience. ACS Chem Neurosci 2, 185–197, doi:10.1021/cn100111a (2011).

54 Zheng, K. et al. Nanoscale diffusion in the synaptic cleft and beyond measured with time-resolved fluorescence anisotropy imaging. Sci Rep 7, 42022, doi:10.1038/srep42022 (2017).

55 Prakash, R. et al. Two-photon optogenetic toolbox for fast inhibition, excitation and bistable modulation. Nat Methods 9, 1171–1179, doi:10.1038/nmeth.2215 (2012).

56 Volterra, A., Liaudet, N. & Savtchouk, I. Astrocyte Ca(2)(+) signalling: an unexpected complexity. Nat Rev Neurosci 15, 327–335, doi:10.1038/nrn3725 (2014).

57 Lohse, M. J., Nuber, S. & Hoffmann, C. Fluorescence/bioluminescence resonance energy transfer techniques to study G-protein-coupled receptor activation and signaling. Pharmacol Rev 64, 299–336, doi:10.1124/pr.110.004309 (2012).

58 Kholodenko, B. N., Hancock, J. F. & Kolch, W. Signalling ballet in space and time. Nat Rev Mol Cell Biol 11, 414–426, doi:10.1038/nrm2901 (2010).

59 Tiwary, P., Limongelli, V., Salvalaglio, M. & Parrinello, M. Kinetics of protein-ligand unbinding: Predicting pathways, rates, and rate-limiting steps. Proc Natl Acad Sci U S A 112, E386–391, doi:10.1073/pnas.1424461112 (2015).

60 Nuñéz, S., Venhorst, J. & Kruse, C. G. Target-drug interactions: first principles and their application to drug discovery. Drug Discov Today 17, 10–22, doi:10.1016/j.drudis.2011.06.013 (2012).

61 Dore, A. S. et al. Structure of class C GPCR metabotropic glutamate receptor 5 transmembrane domain. Nature 511, 557–562, doi:10.1038/nature13396 (2014).

62 Hlavackova, V. et al. Sequential inter- and intrasubunit rearrangements during activation of dimeric metabotropic glutamate receptor 1. Sci Signal 5, ra59, doi:10.1126/scisignal.2002720 (2012).

63 Wijetunge, L. S., Till, S. M., Gillingwater, T. H., Ingham, C. A. & Kind, P. C. mGluR5 regulates glutamate-dependent development of the mouse somatosensory cortex. J Neurosci 28, 13028–13037, doi:10.1523/JNEUROSCI.2600-08.2008 (2008).

64 Morel, L., Higashimori, H., Tolman, M. & Yang, Y. VGluT1+ neuronal glutamatergic signaling regulates postnatal developmental maturation of cortical protoplasmic astroglia. J Neurosci 34, 10950–10962, doi:10.1523/JNEUROSCI.1167-14.2014 (2014).

65 Luscher, C. & Huber, K. M. Group 1 mGluR-dependent synaptic long-term depression: mechanisms and implications for circuitry and disease. Neuron 65, 445–459, doi:10.1016/j.neuron.2010.01.016 (2010).

66 Diering, G. H. et al. Homer1a drives homeostatic scaling-down of excitatory synapses during sleep. Science 355, 511–515, doi:10.1126/science.aai8355 (2017).

67 Panatier, A. et al. Astrocytes are endogenous regulators of basal transmission at central synapses. Cell 146, 785–798, doi:10.1016/j.cell.2011.07.022 (2011).

68 Hagena, H. & Manahan-Vaughan, D. mGlu5 acts as a switch for opposing forms of synaptic plasticity at mossy fiber-CA3 and commissural associational-CA3 synapses. J Neurosci 35, 4999–5006, doi:10.1523/JNEUROSCI.3417-14.2015 (2015).

69 Oh, W. C., Parajuli, L. K. & Zito, K. Heterosynaptic structural plasticity on local dendritic segments of hippocampal CA1 neurons. Cell Rep 10, 162–169, doi:10.1016/j.celrep.2014.12.016 (2015).

70 Jong, Y. I., Harmon, S. K. & O’Malley, K. L. Intracellular GPCRs Play Key Roles in Synaptic Plasticity. ACS Chem Neurosci, doi:10.1021/acschemneuro.7b00516 (2018).

71 Stoppel, L. J. et al. beta-Arrestin2 Couples Metabotropic Glutamate Receptor 5 to Neuronal Protein Synthesis and Is a Potential Target to Treat Fragile X. Cell Rep 18, 2807–2814, doi:10.1016/j.celrep.2017.02.075 (2017).

72 Cabello, N. et al. Metabotropic glutamate type 5, dopamine D2 and adenosine A2a receptors form higher-order oligomers in living cells. J Neurochem 109, 1497–1507, doi:10.1111/j.1471-4159.2009.06078.x (2009).

## Supplemental References

1 Pawlicki, M., Collins, H. A., Denning, R. G. & Anderson, H. L. Two-photon absorption and the design of two-photon dyes. Angew Chem Int Ed Engl 48, 3244–3266, doi:10.1002/anie.200805257 (2009).

2 Gomez-Santacana, X. et al. Illuminating Phenylazopyridines To Photoswitch Metabotropic Glutamate Receptors: From the Flask to the Animals. ACS Cent Sci 3, 81–91, doi:10.1021/acscentsci.6b00353 (2017).

3 Atkinson, P. J. et al. Altered expression of G(q/11alpha) protein shapes mGlu1 and mGlu5 receptor-mediated single cell inositol 1,4,5-trisphosphate and Ca(2+) signaling. Mol Pharmacol 69, 174–184, doi:10.1124/mol.105.014258 (2006).

4 Nash, M. S. et al. Determinants of metabotropic glutamate receptor-5-mediated Ca2+ and inositol 1,4,5-trisphosphate oscillation frequency. Receptor density versus agonist concentration. J Biol Chem 277, 35947–35960, doi:10.1074/jbc.M205622200 (2002).

